# Compact adenine base editors to enable therapeutic rescue of Duchenne muscular dystrophy

**DOI:** 10.64898/2026.05.08.723843

**Authors:** Kartik L. Rallapalli, Joanne M. L. Ho, Isabel Nocedal, Jin Wen Tan, Rodrigo Fregoso Ocampo, Nicole C. Thomas, Brandon Justice, Samatar Jirde, Liliana Gonzalez-Osorio, Cindy J Castelle, Jyun-Liang Lin, Jacquelyn T. Wright, Soungsothira Toch, Krushangi Shah, Benjamin Freeman, Jenat Rahman, Jared Muysson, Isabella Krudop, Lisa M. Alexander, Alan R. Brooks, Christopher T. Brown, Daniela S. Aliaga Goltsman, Kevin G. Hoff, Paul Szymanski, Brian C. Thomas, David W. Taylor, Cristina N. Butterfield

## Abstract

Adenine base editors (ABEs) have emerged as a powerful gene-editing technology enabling precise and programmable adenine-to-guanine substitutions across the genome. However, their translation into in vivo therapeutics is limited by delivery challenges, as their size exceeds the packaging capacity of adeno-associated virus (AAV). Here, we report the discovery, structural characterization, and engineering of two compact, highly active ABEs built on novel deaminases and Cas9d nucleases, enabling all-in-one, single-vector AAV delivery. Applying these compact ABEs to Duchenne muscular dystrophy (DMD), we demonstrate efficient disruption of conserved splice-acceptor sites at dystrophin exons 45 and 51 in human skeletal muscle cells, enabling therapeutically relevant exon skipping. Together, these ABEs help expand the therapeutic reach of base editing towards diverse tissue types and disease targets.

## Introduction

Adenine base editors (ABEs) are an emerging class of gene editing technology that allows the precise and efficient substitution of an A•T base pair to a G•C base pair at any chosen location within the genome **[1]**. These tools offer the ability to directly correct pathogenic single-nucleotide variants (SNVs) or disrupt gene expression without causing double-stranded breaks (DSBs) by creating premature stop codons or alternative splice sites. This capability has already been leveraged towards therapeutically relevant edits for hemoglobinopathies like sickle cell disease **[2, 3]**, alpha-1 antitrypsin deficiency **[4]**, a spectrum of cardiovascular diseases **[5–7]**, cancer immunotherapies **[8]**, and even N-of-1 metabolic disorders **[9]**.

However, the clinical translation of ABE technology has been significantly constrained by delivery limitations. Most current preclinical and clinical base editing efforts have focused primarily on ex vivo applications as well as liver-directed in vivo therapies, due to the substantial size constraints imposed by current delivery vectors. Existing ABE systems often exceed the ∼4.7 kb packaging threshold of adeno-associated virus (AAV) **[10],** necessitating dual-vector split-intein approaches **[11–15]** that increase complexity, require higher viral doses with associated safety concerns **[16]**, and reduce therapeutic efficacy **[17].**

Here, we report on the discovery and engineering of compact AAV-compatible ABEs that unlock the full therapeutic potential of base editing technology across diverse organ systems and tissue types. Starting from novel metagenomically derived adenosine deaminases and compact nucleases, we used cryo-EM structures to design domain-inlaid ABE architectures that demonstrate high editing efficiency while maintaining compatibility with single-AAV delivery. We subsequently applied these compact ABEs to treat Duchenne muscular dystrophy (DMD) by precisely modifying dystrophin exon splice sites, achieving effective exon skipping and dystrophin restoration in human skeletal muscle cells.

## Results

### Discovery and Optimization of Novel Adenosine Deaminases

An ABE mechanistically functions as a two-component system: a programmable CRISPR nuclease mediates the DNA targeting and R-loop formation, while an adenosine deaminase (ADA) catalyzes the conversion of desired adenosine to inosine within the exposed single-stranded DNA. Endogenous repair pathways subsequently process inosine as guanine, resulting in a permanent A→G transition. Thus, fusion of deaminase with a nicking nuclease generates an ABE capable of directing precise A•T-to-G•C edits at any pre-defined genomic locus **[1].**

To create ABEs compatible with single-AAV delivery, we capitalized on the intrinsic modularity of the ABE architecture and redesigned the platform around compact nuclease and deaminase elements rather than relying on *Streptococcus pyogenes* Cas9 (*Sp*Cas9) nuclease and the *Escherichia coli* TadA (*Ec*TadA) deaminase.

Previously, we reported the discovery, characterization, and optimization of a novel class of Type II-D nucleases that are approximately half the size of conventional Type II-A systems such as *Sp*Cas9 **[18].** Here, we prioritized two novel compact Cas9d nucleases: MG34-29 (747 amino acids) and MG102-71(943 amino acids), where “MG” denotes their metagenomic origin **(Fig. 1A)**. As described previously, MG34-29 is an ancestrally reconstructed nuclease derived from the Cas9d MG34 clade and exhibits 12-13-fold higher activity than its modern descendants **[19].** Given the successful results observed with the MG34 family of nucleases, we extended the use of ancestral sequence reconstruction (ASR) methods to the MG102 family of nucleases for protein sequence diversification and activity engineering **[18, 19]**.

**Fig. 1:**
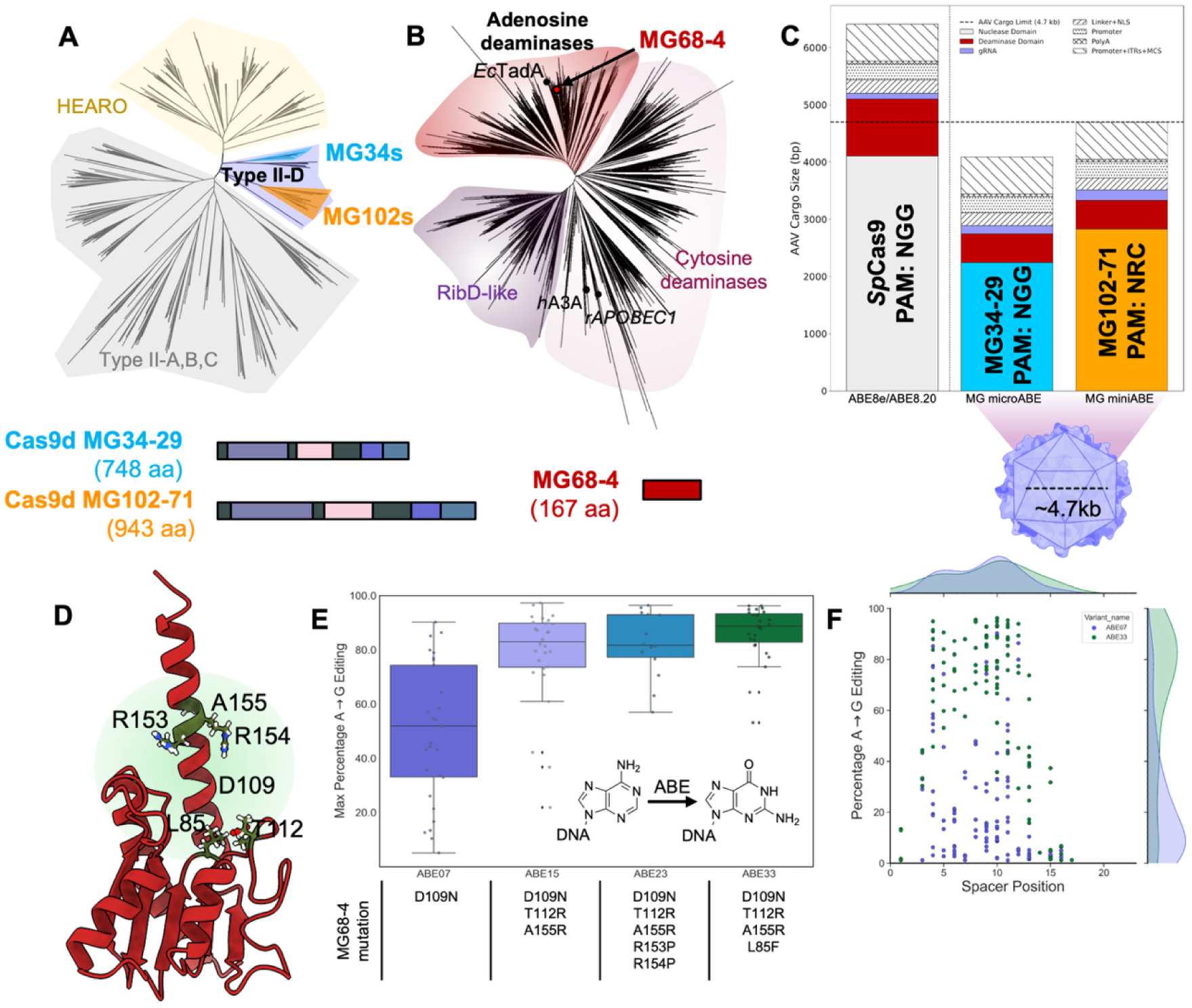
Identification and engineering of a compact adenosine deaminase for AAV-compatible adenine base editors. **(A)** Phylogenetic tree of Type II CRISPR nucleases, highlighting compact Type II-D nucleases relative to canonical Type II-A Cas9 clade; schematic size comparisons are shown below. **(B)** Phylogenetic tree of deaminases, highlighting the MG68-4 forming a distinct clade from cytidine deaminases. **(C)** Size comparison of *Sp*Cas9-, MG34-29-, and MG102-71-based ABEs demonstrating compatibility of Cas9d ABEs with single-AAV packaging. **(D)** AlphaFold-predicted structure of MG68-4 with engineered residues proximal to the active site highlighted. **(E)** Max A→G editing efficiency of MG68-4 variants across a panel of 15 distinct genomic sites at two loci. Boxes indicate median and interquartile range; whiskers denote full range. **(F)** Positional editing profiles across 15 guides at two loci showing a broad MG68-4 editing window spanning protospacer positions 2-15, largely conserved across deaminase variants ABE07 and ABE33.

In this study, we generated MG102-71 by reconstructing the MG102 clade **(Supplementary Note 1)**. MG102-71 shares 76.5% sequence identity with MG102-2 **(Fig. S1A, S1B)**, its closest extant relative, and demonstrates up to two-fold increase in overall nuclease activity in mammalian cells. Notably, MG102-71 is also active across more targets relative to representative modern clade members **(Fig. S1C).**

MG34-29 recognizes an NGG PAM **[18, 19]**, whereas MG102-71 recognizes an NRC PAM **(Fig. S1D)** (where N denotes any nucleotide and R denotes a purine). Together, these two compact nucleases not only enable targeting of sites accessible to canonical *Sp*Cas9 but also expand access to additional genomic loci, increasing the overall targetable genomic space of compact ABEs **(Fig. 1)**.

Beyond simply miniaturizing the nuclease component, we identified an additional opportunity to reduce the size of the deaminase component. The *Ec*TadA used in canonical ABE functions as an obligate dimer, with standard ABE architectures employing both a wild-type and an evolved TadA* monomer **[1, 20–22]**, effectively doubling the deaminase footprint to ∼300 amino acids. To overcome this constraint, we sought compact adenosine deaminases that could operate as independent monomers, thereby enabling further reduction in the total ABE size.

To identify novel deaminases compatible with our compact nuclease scaffolds, we mined a large database of high-quality, assembled microbial metagenomes from diverse environments, yielding thousands of candidate deaminase genes. Novel families and subfamilies were defined and annotated using *Ec*TadA and well-characterized cytosine deaminases, rat APOBEC1 and human APOBEC3A, which serve as the catalytic domains of widely used cytosine base editors. Candidate ADAs were nominated based on sequence and structural hallmarks, such as the presence of the conserved catalytic -CxxC- and -HxE- motifs. Selected candidates were next evaluated as base editors using a bacterial selection assay **(Fig. S2, Supplementary Note 2)**.

From this search, we identified a promising candidate, designated MG68-4, where “MG” once again signifies its metagenomic discovery. MG68-4 shares less than 70% average amino acid identity with *Ec*TadA and forms a distinct phylogenetic clade close to well-characterized deaminases **(Fig. 1B)**. Based on this phylogenetic separation and its homology to TadA, we hypothesized that MG68-4 functions as an ADA.

Given the modular nature of ABEs and the limited functional information available for MG68-4, we first sought to characterize and optimize MG68-4 within the cognate ADA dimeric architecture fused within our previously described Type II-A *Sp*Cas9-like nuclease, MG3-6_3-8 **[23, 24] (Supplementary Note 2)**. We rationally engineered MG68-4 for enhanced catalytic performance **(Fig. 1D)**. Accordingly, we targeted twelve residues in or near the predicted MG68-4 active site **(Fig. S3, S4).** We evaluated the performance of the MG68-4 ABE variants across a panel of 15 gRNAs targeting *mApoA1* and *mAngptl3* with a diverse variety of A positions and A neighbors.

We initiated engineering of MG68-4 by introducing D109N, the first mutation evolved in *Ec*TadA during the ABE7.10 campaign **[1],** which is well-characterized to enhance DNA binding of EcTadA directly **[20, 25].** This initial variant, termed ABE07, exhibited only ∼11% mean A→G editing across all adenines within the 15-guide panel.

To improve activity, we next incorporated two highly impactful mutations from later ABE evolution efforts: T112R, developed in the ABE8e campaign **[26]**, and A155R, which resides on the α5 helix of the deaminase. Introducing these two mutations simultaneously into ABE07 produced a 1.7-fold increase in mean editing, yielding the variant ABE15.

Upon evaluating additional mutations on the ABE15 background, we identified two high-performing variants: ABE23, containing R153P and R154P adjacent to A155R, and ABE33, containing L85F within the deamination active site. Notably, although these mutations significantly elevated mean editing efficiency across diverse genomic contexts, the maximum editing observed at preferred adenines remained similar to that of ABE15 **(Fig. 1E)**. This finding suggests that MG68-4 possesses intrinsically high catalytic potential on optimal substrates, and that mutations beyond the core set, D109N, T112R, and A155R, may demonstrate catalytic benefits in a context-dependent manner.

A second key observation from our engineering efforts is that the editing window of the ADA domain in MG68-4 spans positions 2-15 of the exposed R-loop. This window remained essentially unchanged between the minimally evolved ABE07 and the highly optimized ABE33 **(Fig. 1F).** These results indicate that the MG68-4 deaminase naturally acts on any adenine exposed within the R-loop and that our engineering did not alter the inherent breadth of the editing window.

Overall, these results show that a novel putative ADA, MG68-4, can be transformed into a highly efficient ABE through only a few strategic point mutations, achieving near-saturating A→G editing across diverse genomic contexts.

### Structure-guided engineering of compact ABEs with enhanced activity

Previous studies **[27–30]**, including our own work **(Supplementary Note 2)**, have demonstrated that inlaying the deaminase domain within the nuclease chassis, especially Type II-As, facilitates productive R-loop engagement and activates the base editing activity of the resultant ABE. To extend this strategy toward the creation of compact ABEs, we first sought to obtain a high-quality structure of MG34-29 in its R-loop-bound conformation using cryo-EM.

We expressed MG34-29 in *E. coli* to homogeneity and determined the cryo-EM structure of the MG34-29-gRNA-target DNA complex with a resolution of 3.1 Å **(Fig. 2A)**. Consistent with our previous structural studies, MG34-29 adopts an overall bilobed architecture similar to that observed for MG34-1 **[19]**, with an overall Cα RMSD of 3.4 Å between the nuclease structures and 2.9 Å between the phosphate backbone of the gRNA scaffolds. The most notable structural differences emerge in the RuvC and REC domains. The RuvC domain, which directly contacts the R-loop, is substantially better resolved in the MG34-29 structure compared to MG34-1. This improved resolution likely reflects enhanced stabilization of the R-loop-bound state and is consistent with the higher nuclease activity observed for MG34-29. In both structures, the HNH domain is unresolved, suggesting that this domain remains highly flexible post-cleavage for both effectors. Intriguingly, a significant portion of the REC domains in MG34-29 is poorly resolved, unlike the MG34-1 model, indicating that the REC domain of the ancestrally reconstructed MG34-29 nuclease may exhibit suboptimal compatibility with the native MG34-1 gRNA scaffold.

**Fig. 2:**
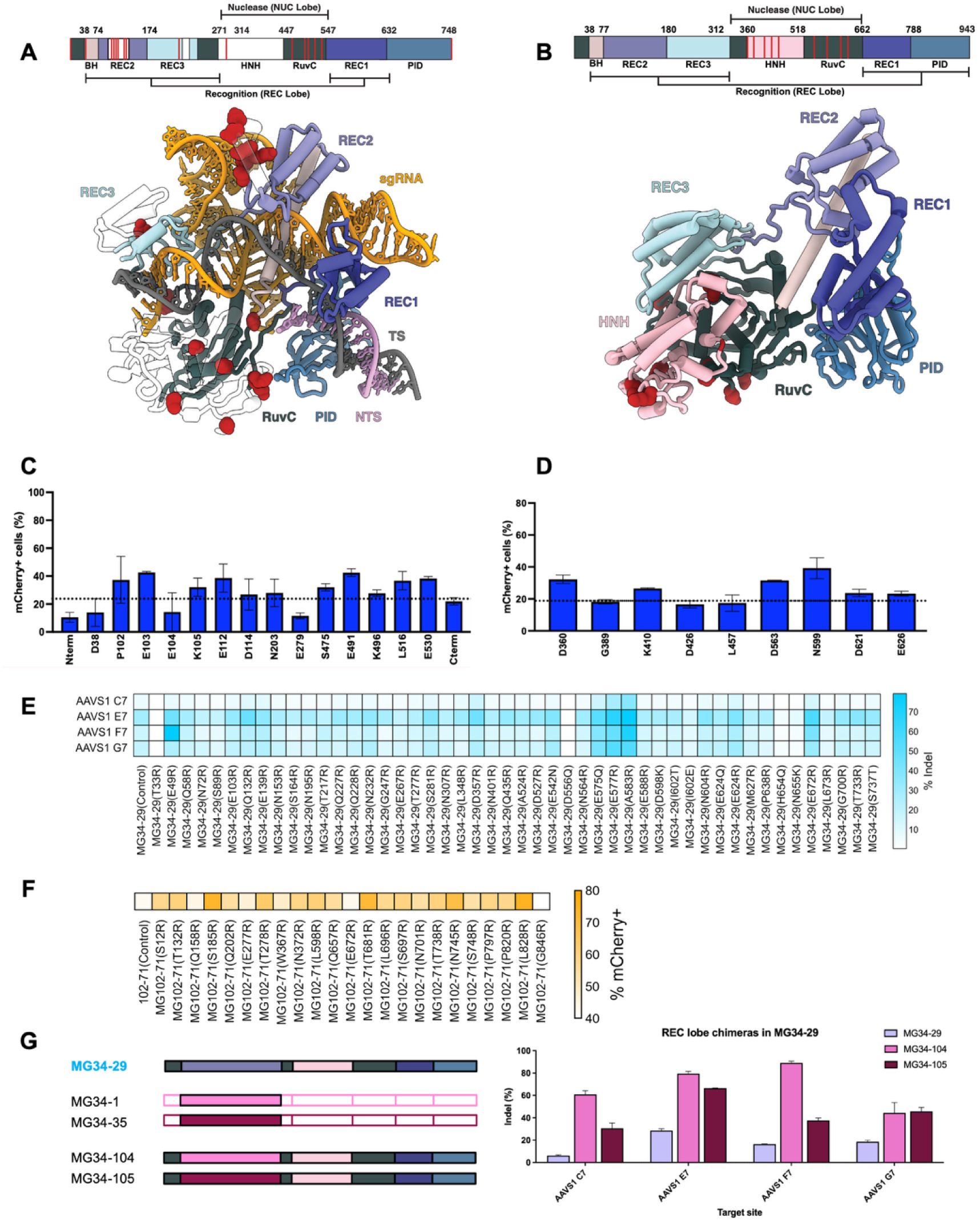
Structure-guided engineering of compact Type II-D nucleases to affect base editing activity. Structure of **(A)** MG34-29 resolved using cryo-EM and modelled using Boltz, and **(B)** MG102-71 predicted using Boltz, highlighting domain architecture and selected inlaid insertion sites for MG68-4 deaminase domain. Editing activity of compact ABEs created by inserting MG68-4 engineered deaminase within **(C)** MG34-29 and **(D)** MG102-71, quantified using a mCherry-based reporter assay in HEK293T cells and normalized to nuclease-only controls. **(E)** Editing efficiencies (indels) for MG34-29 arginine scanning variants across multiple AAVS1 target sites. **(F)** Editing efficiencies (%mCherry turn on) MG102-71 ABEs at multiple spacers. **(G)** REC-lobe chimeragenesis in MG34-29. Metagenomically mined REC-domain swaps generate chimeric compact ABEs with substantially enhanced editing activity across multiple targets. Bars represent mean ± s.d.; n = 2 biologically independent replicates.

We leveraged this cryo-EM structure together with recent advancements in AI-structure prediction algorithms to generate full-length structural models of both MG34-29 and MG102-71 nucleases **(Fig. 2A, B) [30]**. Guided by these structural models, we identified solvent-exposed loops within the flexible regions of MG34-29 and MG102-71 as candidate sites to inlay the engineered MG68-4 deaminases. These variants were cloned into a mCherry reporter cell line that quantitatively reports targeted A→G editing **(Supplementary Note 3, Figure S4) [28, 31]**.

For MG34-29, inlaying the deaminase domain consistently resulted in editing efficiencies that were equal to or greater than those observed with C-terminal fusion; interestingly, the N-terminal fusion showed the lowest activity. The highest A→G editing activity was observed for inlay positions E103 and E112 within the REC domain, and E491 and L516 within the RuvC domain, which exhibited approximately three- to four-fold higher activity relative to the canonical N-terminal fusion architecture **(Fig. 2C)**. Guided by these results from MG34-29, we next focused on the analogous endonuclease domains of MG102-71 and restricted deaminase inlay to the RuvC and HNH regions. Within this framework, inlay of the engineered MG68-4 deaminase at position N599, once again located in the RuvC domain of MG102-71, yielded the highest editing activity among the tested configurations **(Fig. 2D)**. Intriguingly, the highest performing insertion sites correspond to the unresolved regions of the MG34-29 cryo-EM structure, suggesting that targeting the conformationally labile or unresolved regions can be used to successfully insert functional domains within any nuclease chassis **(Fig. 2A)**. This observation that the RuvC domain of these novel Type II-D nucleases is particularly amenable to deaminase inlay is consistent with reported functional inlaid base editors based on other Type II-A nucleases, including SpCas9 **[27]**, Nme2Cas9 **[28]**, SaCas9 **[29]**, and MG3-6_3-8 **(Supplementary Note 2).**

In parallel, we further leveraged these structural models to rationally engineer variants that increase positive electrostatic potential on the nuclease surface and the regions in close proximity to each effectors’ nucleic-acid binding partners. We hypothesized that introducing arginine or polar amino acids would strengthen nucleic acid interactions, thereby improving both the nuclease activity and base editing efficiency. For MG34-29, this approach revealed several gain-of-function variants, with substitutions at E49R, E577R, and A583R producing the largest improvements **(Fig. 2E).** The E49R mutation, located in the bridge helix, yields approximately 2.2-fold increased activity, likely through the formation of new non-specific electrostatic interactions with the gRNA and DNA in the PAM-proximal region. Importantly, this interaction occurs with the phosphate backbone rather than the nucleobase, suggesting that the enhancement in activity is sequence independent. The E577R and A583R, located within the REC1 domain, conferred approximately 2.8-fold and 3.4-fold enhancements, respectively, and could be explained by introducing stabilizing ionic interactions with the target strand DNA upstream of the PAM.

Similarly, arginine substitutions in MG102-71 produced several high-performing variants. Among these, L828R, S185R, and T681R displayed the strongest effects, with 1.77-fold, 1.73-fold, and 1.69-fold increases in editing activity, respectively, compared with the MG102-71 control **(Fig. 2F)**.

Finally, our cryo-EM structure of MG34-29 revealed a substantial unresolved portion of the REC domain, unlike that of the native MG34-1 structure **(Fig. 2A)**, consistent with our previous observation that the sequence conservation within this region is relatively low, with MG34-29 sharing only 71% sequence identity with MG34-1 **[19]**. Because the REC domain plays a critical role in gRNA binding, we hypothesized that the REC lobe of ancestrally reconstructed MG34-29 may be suboptimally compatible with the native MG34-1 gRNA scaffold. To test this hypothesis, we performed REC-lobe chimeragenesis, replacing the REC domain of MG34-29 with corresponding REC lobes derived from the naturally occurring MG34-1 and its closely related naturally occurring homolog MG34-35. The resulting chimeric nucleases, MG34-104 and MG34-105, exhibited substantially improved activity when paired with the native MG34-1 guide RNA scaffold compared with MG34-29 **(Fig. 2G)**.

Building on the individual engineering strategies described above, we combined the top-performing elements identified from deaminase optimization **(Fig. 1),** structure-guided deaminase inlay **(Fig. 2C, D),** arginine scanning **(Fig. 2E, F),** and REC-lobe chimeragenesis **(Fig. 2G)** into unified compact ABE architectures. The resulting editors were evaluated for A→G editing activity in HEK293T cells across multiple AAVS1 safe-harbor target sites.

For MG34-29–based ABEs, hereafter designated as microABEs, integration of the optimized MG68-4 variant with REC(E112) or RuvC(L516) deaminase inlay and arginine substitutions yielded the highest-performing constructs, with microABE267 consistently exhibiting the greatest A→G editing activity across tested loci, reaching approximately 60% total A→G conversion **(Fig. 3A)**. In contrast, MG34-29 editors lacking REC-lobe chimeragenesis or arginine scanning exhibited more modest activity, underscoring the additive effects of these engineering strategies. To understand the structural impact of these modifications, we determined the cryo-EM structure of microABE273, which incorporates the largest number of arginine substitutions among the MG34-29 variants **(Fig. S8)**. In this structure, although the MG68-4 deaminase domain remained unresolved, its fusion within the RuvC domain led to the stabilization and resolution of the elusive HNH domain.

**Fig. 3.**
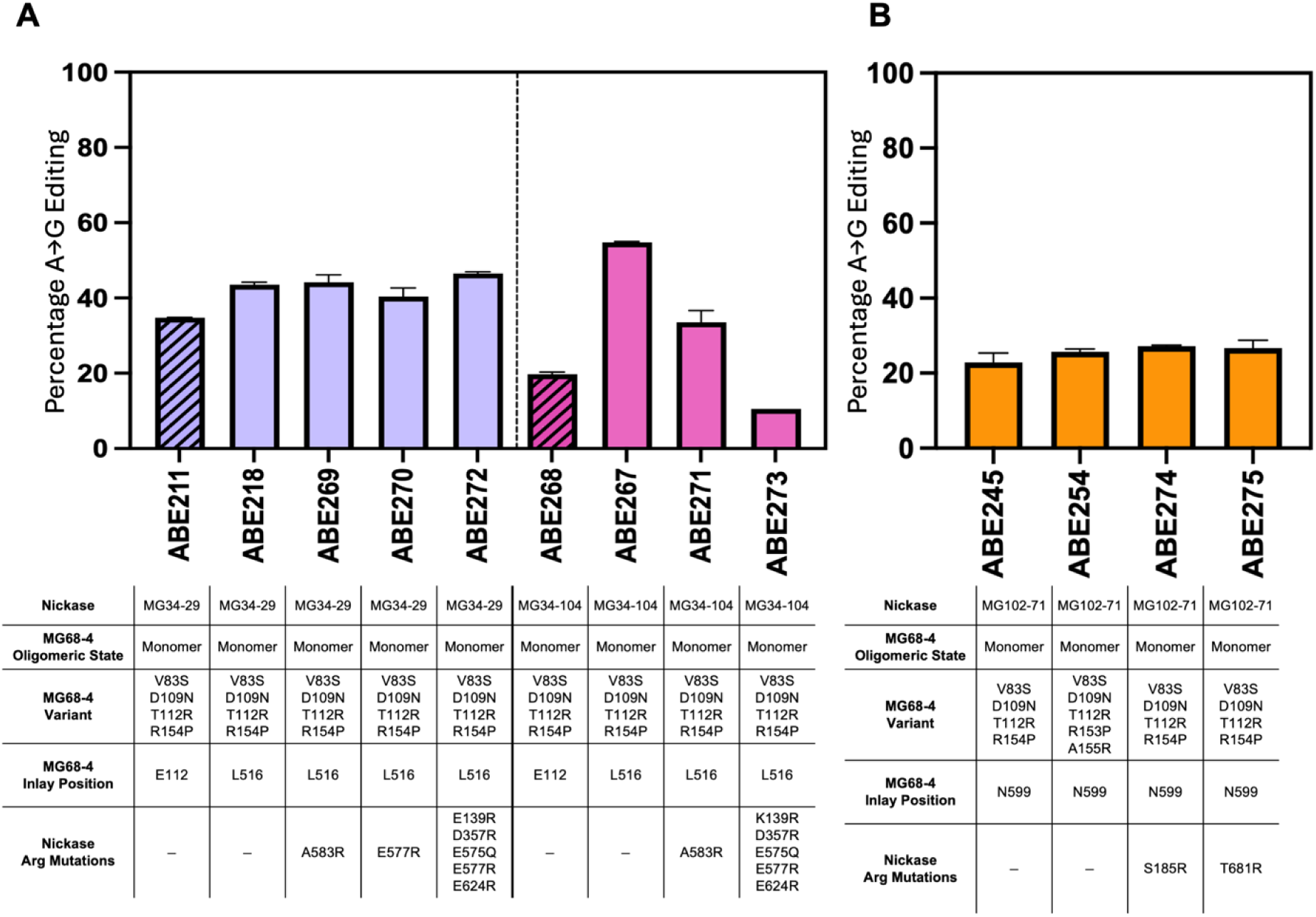
Combinatorial engineering of nuclease and deaminase to establish the best compact ABE. Maximum A→G editing frequencies for **(A)** MG34-29 microABEs and **(B)** MG102-71 miniABEs incorporating different MG68-4 variants, inlay positions, and nickase variants in SkM cells. Editor configurations and key mutations are annotated below each bar. Bars represent mean ± s.d.; *n*=2 biologically independent replicates.

Similarly, for MG102-71-based ABEs, hereafter designated as miniABEs, combining the MG68-4 deaminase inlay at the N599 within RuvC and arginine substitutions produced the most active miniABE variants. Specifically, miniABE275 showed the highest activity, achieving 35-40% A→G editing across the AAVS1 target site **(Fig. 3B)**. Together, these results demonstrate that systematic integration of complementary engineering strategies can yield compact ABEs with substantially enhanced activity while remaining compatible with single-AAV delivery.

### Compact ABEs Can Enable Therapeutic Exon Skipping to Rescue DMD

Having established the efficiency of our compact ABEs, we next evaluated their performance in a therapeutically relevant context by targeting Duchenne muscular dystrophy (DMD), a severe genetic disorder caused by frameshifting or nonsense mutations in the DMD gene that abolishes dystrophin expression, leading to progressive muscle degeneration and early mortality **[32]**. We focused our targeting strategy on the mutational hotspot spanning exons 45-55 of the DMD gene, a region in which deletions frequently disrupt the dystrophin reading frame and produce an internally truncated yet functional dystrophin protein **(Fig. 4A) [33, 34]**.

**Fig 4:**
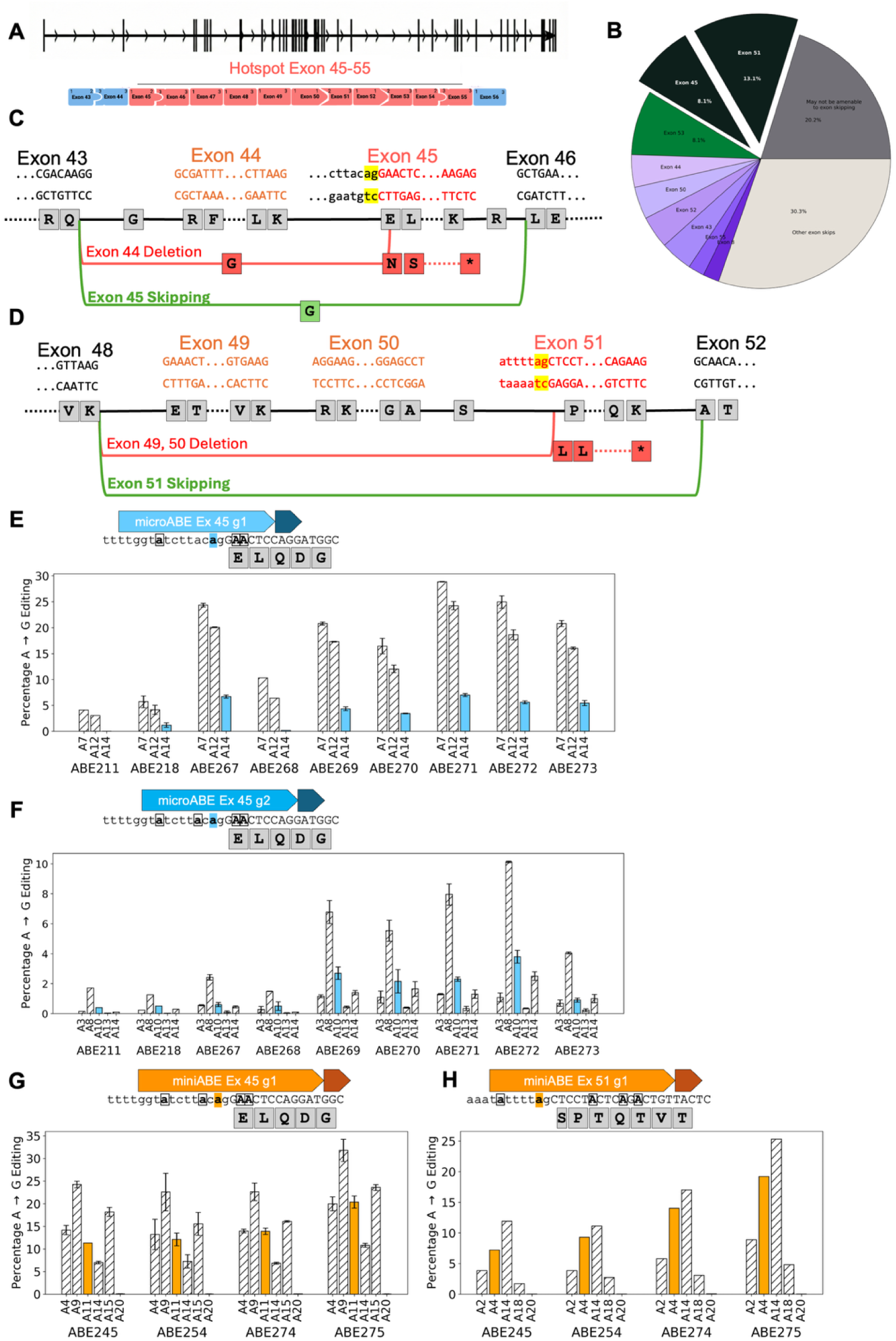
Compact ABEs to enable splice-site disruption and exon skipping in the DMD gene. **(A)** Schematic of the DMD locus highlighting the mutation hotspot spanning exons 45-55. **(B)** Distribution of DMD patient mutations amenable to exon-skipping interventions across candidate exons. **(C)** DMD patients with an exon 44 deletion exhibit a disrupted reading frame (red) compared to wild-type DMD (black). An A→G conversion at the conserved splice acceptor adenine of exon 45 induces exon 45 skipping, restoring the reading frame despite the exon 44 deletion (green). **(D)** DMD patients with an exon 49 and/or 50 deletion exhibit a disrupted reading frame (red) compared to wild-type DMD (black). An A→G conversion at the conserved splice acceptor adenine of exon 51 induces exon skipping, restoring the reading frame despite these deletions (green). Bar plots showing A→G editing efficiencies at DMD exon 45 **(E-G)** and exon 51 **(H)**. The target splice-site adenine is highlighted with solid bars (blue, microABEs; orange, miniABEs), while bystander adenine editing is indicated by hashed bars. Editing efficiencies are presented as mean values, and error bars represent the standard deviation of two biological replicates.

Skipping of exon 45 is predicted to benefit patients carrying common upstream deletions that disrupt the reading frame, most frequently deletions involving exon 44, by re-establishing an in-frame transcript through the removal of exon 45. This strategy is estimated to apply to approximately 8.1% of individuals with Duchenne muscular dystrophy **(Fig. 4B, C)**. Skipping of exon 51 targets a complementary patient population harboring frequent downstream multi-exon deletions within the hotspot locus, and is predicted to benefit approximately 13.1% of patients with DMD **(Fig. 4B, D)**. Indeed, multiple therapeutic strategies based on exon 45 or exon 51 skipping have been clinically validated with several antisense oligonucleotide therapies for the treatment of DMD **[35–37]**.

To explore the applicability of our compact ABEs towards treating DMD, we prioritized these high-impact targets. We designed gRNAs positioning the conserved splice acceptor adenines of exons 45 and 51 within the active deamination window of our ABEs and tested them in human skeletal muscle (SkM) cells, a physiologically relevant cell type for DMD. Robust editing activity was observed across multiple ABEs, with the target splice-site A accounting for a substantial proportion of total A→G edits **(Fig. 4E-H)**.

At exon 45, the miniABE275 exhibited the highest activity, achieving splice-site A→G editing of 20.3% and 34.6% total cumulative A→G editing across all editable adenines within the protospacer sequence. In contrast, the microABE271 showed comparable total protospacer-wide A→G editing (31.6%) but reduced splice-site–specific editing (7.0%), highlighting differences in editing specificity between editor nickase chassis. Editing efficiencies were also guide-dependent, with microABE gRNA g1 exhibiting approximately three-fold higher activity than gRNA g2 **(Fig. 4F, H)**.

For exon 51, only miniABE variants could be evaluated owing to their more permissive NRC PAM requirement. Consistent with exon 45 results, miniABE275 again demonstrated the highest activity, achieving splice-site A→G editing of 19.2% and total A→G editing of 28.5% **(Fig. 4H)**.

Across the tested variants, we observed differences in on-target A and bystander substitution profiles, which were consistent with the broad editing windows characterized in **Fig. 1F**. Because the splice-acceptor function is determined by extended sequence motifs beyond the invariant AG dinucleotide, these bystander edits near the splice acceptor motif may further enhance exon skipping and increase the yield of truncated yet functional dystrophin gene product. Together, these initial findings establish the feasibility of compact ABEs for AAV-based gene editing strategies.

## Discussion

Previous studies have sought to generate AAV-packageable ABEs through split-intein strategies, truncation of *Sp*Cas9 nuclease domains **[38]**, use of a single TadA copy **[27]**, the use of smaller Cas9 orthologs **[31, 39, 40],** and, in some cases, compact Cas12-based editors **[41–44]**. More recently, alternative programmable systems such as IscB transposon-associated nucleases **[45–47]** and TnpB endonucleases **[48]** have been explored as potential chassis for base editing. For example, ABE8e-mediated editing of splice acceptor A→G at exon 45 and exon 51 has been shown to successfully restore therapeutic levels of dystrophin mRNA and protein in mouse models, albeit requiring a dual-AAV split delivery system **[49, 50]**.

The miniABEs and microABEs developed in this study demonstrate high base-editing efficiency while remaining within the packaging limits of AAV vectors, enabling up to 30% productive editing at splice acceptor sites in human skeletal muscle cells. These results provide a foundation for single-administration single-AAV therapeutic strategies that could apply to a substantial subset of patients with DMD. More broadly, these compact ABEs may expand the therapeutic reach of base editing across a range of extra-hepatic in vivo targets.

## Materials & Methods

### Computational Enzyme Discovery and Structure Analyses

4 Tbp of proprietary or public assembled metagenomic sequencing data from diverse environments (soil, sediments, groundwater, thermophilic, human, and non-human microbiomes) were mined to discover novel deaminases. HMM profiles of known deaminases were built and searched against all predicted proteins using HMMER3 (hmmer.org) to identify all deaminases from our databases. Predicted and reference deaminases were aligned with MAFFT **[51, 52]**, and a phylogenetic tree was inferred using FastTree2 **[53]**. Candidate deaminases were prioritized based on the presence of conserved catalytic residues and key structural motifs characteristic of reference adenosine deaminases.

### Cloning, expression, and purification

The *E. coli* codon-optimized DNA sequence of MG34-29 was cloned in the pET21(+) backbone using Gibson assembly methods. The MG34-29-pET21(+) expression vector was transformed into *E. coli* BL21-AI (Invitrogen) chemically competent cells and overexpressed in 2L bioreactors. Cells were grown at 30°C until the desired OD of 20 was reached. Cultures were subsequently induced with 0.22% Arabinose and 0.2 mM IPTG at 25°C and grown for 18 hours before harvesting. 75 g of total cell paste was thawed and resuspended in Buffer A (50 mM CHES pH 9.0, 250 mM NaCl, 1 mM MgCl2, 0.5 mM EDTA, 20 mM Imidazole, 10% glycerol) supplemented with Pierce Protease Inhibitor Tablets, lysozyme and 0.5% n-octyl-β-d-glucoside. Cells were lysed by sonication, and the lysate was clarified by centrifugation at 35,000xg for 30 minutes. Cell lysate supernatant was loaded onto a 5 mL HisTrap HP Ni Column (Cytiva) and eluted using a gradient of Buffer B (50 mM CHES pH 9.0, 250 mM NaCl, 1 mM MgCl2, 5% glycerol) and elution buffer (buffer C: 50 mM CHES pH 9.0, 250 mM NaCl, 1 mM MgCl_2_, 0.5 mM EDTA, 500 mM Imidazole, 10% glycerol). Fractions containing the protein were combined and injected onto a HiTrapQ FF (Cytiva) anion exchange chromatography column and eluted with buffer C (50 mM HEPES pH 7.5, 2 M NaCl, 5 mM MgCl_2_, 0.5 mM EDTA, 5% glycerol). Fractions with A280 > A260 were pooled, concentrated and further purified using a S200i (Cytiva) size exclusion chromatography column using Buffer B as the eluent. Fractions with A280 > A260 were pooled, concentrated, aliquoted and snap frozen in LN2 and stored at -80°C.

The MG34-104 effector was cloned into a pET28 expression vector containing a T7 promoter under lacO repression with a 6x N-terminal His tag and a TEV protease site. A pBR322 vector containing the full 158bp sgRNA under the expression of an E. coli constitutive ribosomal promoter J23119 with a poly-T region directly downstream of the sgRNA sequence for transcription termination was co-transformed with the 34-104 pET28 plasmid into OverExpress C41 (DE3) (Sigma-Aldrich) chemically competent cells. A single colony was inoculated into 50 mL of Luria Broth (LB) under kanamycin and chloramphenicol antibiotic selection and was allowed to grow at 37 °C for 16 h. The inoculate was then back diluted 1:100-fold into two separate 1.5 L LB culture flasks supplemented with a final concentration of 50uM ZnCl2 (TFS) and grown to OD600 0.8. Expression was induced by addition of

Isopropyl-β-D-thiogalactopyranoside (IPTG) at a final concentration of 0.1 mM and growth for 16 h at 18 °C to maximize expression levels. Cells were pelleted by centrifugation (4000 × g, 60 min) and resuspended using Lysis Buffer (20 mM tris-HCl pH = 7.5, 500 mM NaCl, 10 mM MgCl2, 1 mM TCEP, 200 μM PMSF, 10% glycerol) with 1 Pierce Protease Inhibitor Tablet and 10U DNAse I. Cells were lysed by sonication and the lysate was clarified by centrifugation at 18,000 × g for 45 min at 4 °C. The supernatant was loaded into a 5 mL HisTrap HP Ni Column (Cytiva) and eluted using a gradient of Ni elution buffer (20 mM tris-HCl pH = 8.0, 500 mM NaCl, 250 mM Imidazole, 1 mM TCEP, 10% glycerol). Fractions containing the protein were combined and dialyzed with TEV protease in dialysis buffer (40 mM HEPES pH = 8.0, 150 mM NaCl, 1 mM TCEP, 10% glycerol) for 16 h at 4 °C. The sample was then purified by gel filtration chromatography (Superose 6 10/300; GE Healthcare) in dialysis buffer (150 mM NaCl, 200 mM KCl, 1 mM EDTA, 20 mM HEPES). Purity and quality of the protein were analyzed by SDS-polyacrylamide gels. The protein was quantified using the PIERCE Bradford assay reagent kit (Thermo Fisher) following the manufacturer’s instructions.

### Cryo-EM sample preparation, data collection, and processing

Prior to complex assembly, a single aliquot of 50μM sgRNA (IDT) was heated at 95 °C for 2 min followed by cooling at 0.2°C/s to room temperature in 20mM HEPES buffer pH = 7.5. The target dsDNA duplex was formed by mixing equimolar ratio of target and non-target strand, heated at 95 °C for 5 min, then slow cooling at 0.2°C/s to room temperature in 20mM HEPES buffer pH = 7.5. For the MG34-29 complex, a flash frozen aliquot of purified and concentrated MG34-29 was rapidly thawed and incubated at a 10uM final concentration using a 1:1.2:1.2 enzyme:target DNA:sgRNA ratio at 37 °C for 1 h. For the MG34-104 structure an 8-azanelaburine was included in the NTS at the 12^th^ nucleotide of the spacer. All samples were applied to Quantifoil 1.2/1.3 400-Mesh Carbon Grids that had been glow-discharged with 20 mA for 30 s. All grids were blotted using a Vitrobot Mark IV (Thermo Fisher) for 7-10s, blot force 0 at 4 °C and 100% humidity, and plunge-frozen in liquid ethane.

For the MG34-29 complex, data were collected on an FEI Titan Krios 300 V Transmission Electron Microscope (Thermo Scientific) with a Gatan BioContinuum Imaging Filter and a K3 direct electron detector. Data were collected on Serial EM v3.9 with a 0.832 Å pixel size and a defocus range of –1.5 to –2.5 μm with a total exposure time of 6 s in a total accumulated dose of 70e/A^2^. Reference-based motion correction, contrast transfer function (CTF) estimation and particle picking were carried out in cryoSPARC live v4.7. A total of 4,581 movies were accepted at –30° tilt. For the MG34-104 structure,1686 movies were accepted using an FEI Glacios cryo-TEM equipped with a Falcon 4 detector with a pixel size of 0.933 Å. Data for the 34-104 complex was not collected at tilt. All subsequent data was processed using cryoSPARC v4.7 For the MG34-29 complex, 5 8,030,324 particles were selected using template picker using a previously solved Cas9d structure (PDB: 9AUF) **[19]**. A total of 2,834,933 particles were picked after a single round of 2D classification. A random subset of 100,000 particles were picked for *ab-initio* reconstruction (3 classes). The best class was used for heterogeneous refinement and downstream non-uniform refinement, yielding a 2.85 Å reconstruction. These particles were subsequently subclassified using 3D classification. Homogenous particles were chosen for the final 3.1 Å reconstruction comprising 282,286 particles. For the MG34-104 complex 1,558,277 particles were selected using the same methods outlined for the 34-29 complex. Picked particles were subjected to a single round of 2D classification (50 classes). A random subset of 100,000 particles were picked for *ab-initio* reconstruction (3 classes). After ab-initio reconstruction the best class was used along with 582,771 particles from 2D classification for heterogeneous refinement and downstream non-uniform refinement, yielding a 3.34 reconstruction. Subsequent 3D classification was performed for this structure but no deaminase domains were present in any of the classes.

### Cryo-EM model building and structural refinement

The initial model for 34-29 was built by rigid body fitting previously solved structures of Cas9d (PDB: 9AUF) **[19]**. This structure displayed very similar characteristics to previously solved Cas9 structures. This model was used as a fiducial for de-novo model building using a combination of Coot v1.0 **[54]** and ChimeraX v1.90 **[55].** REC3 domains were deleted from the complex, and an additional region of the RuvC domain was built de novo from using the previous template using a combination of AlphaFold **[56]**, rigid body fitting and Coot v0.9.8.91 **[54]**. Multiple iterations of real space refinement implemented within Phenix v1.20.1 were carried out to improve the model geometry and map fit **[57]**. The 34-104 model was built using Cosmic2 ModelAngelo **[58]**. The model was subsequently manually refined using COOT v0.9.8.91 and Isolde. Multiple iterations of real space refinement were also carried out within Phenix v1.20.1 until a final model was reached. Three-dimensional models supporting deaminase and nuclease engineering were predicted using Boltz-2 **[30]** and analyzed using UCSF ChimeraX v1.90 **[55]**.

### Molecular Biology

Sequences for nucleases and ABEs with 5’ and 3’ UTRs were codon optimized for human expression and each cloned into an expression vector either with or without a CleanCap T7 promoter and a polyA tail (Twist Biosciences). The coding sequence contained an N-terminal SV40 nuclear localization signal and a C-terminal nucleoplasmin nuclear localization signal. If not present in the expression vector, a CleanCap T7 promoter and the polyA tail were added via PCR amplification. Individual point mutants were generated via site-directed mutagenesis using the NEBuilder HiFi DNA assembly kit and confirmed via whole plasmid sequencing. The expression vector was midi-prepped (Zymo Research), linearized with SpeI (NEB), cleaned with SureClean (Meridian Bioscience), and used for in vitro transcription with Hi-T7 (NEB). PCR amplified nuclease coding sequences were directly used for in vitro transcription with Hi-T7 RNA polymerase (NEB). In vitro transcription reactions contained N1-methylpseudouridine in place of uridine and had added CleanCap reagent (TriLink Biotechnologies). The resulting mRNA was cleaned with RNeasy (Qiagen), checked for product size and purity by NanoDrop and Tapestation (Agilent) and diluted to 250 ng/uL in sterile water for use in nucleofection.

Combinatorial mutants were generated from two piece NEBuilder cloning of single mutant backbones and gene fragments containing the mutations of interest, then prepared as mRNA following the same procedure.

The base editing efficiency of these small ABEs was assessed across target guides shown in **Table S3.** The first and last three bases on these guides were modified with 2’-O methyl and phosphorothioate groups.

### Mammalian Cell Culture and Transfections

Hepa1-6 (ATCC #CRL-1830) and K562 (ATCC #CCL-243) cells were cultured in IMDM + GlutaMAX (Corning) supplemented with 10% FBS (Corning) for 1–2 passages prior to nucleofection. Cells were harvested, washed with 1× PBS, and resuspended in SF buffer (Lonza Technologies) according to the manufacturer’s instructions. For each condition, 120,000 cells were nucleofected with 500 ng mRNA and 150 pmol sgRNA (for Hepa1-6) or 200 pmol (for K562) using the SF Cell Line 96-well Nucleofector™ Kit (Lonza Technologies). SkM cells (ATCC #PCS-950-010) were maintained in Mesenchymal Stem Cell Basal Medium supplemented with the Primary Skeletal Cell Muscle Growth Kit and seeded at 12,500 viable cells per well in TC-treated 96-well plates (200 µL per well) one day prior to transfection. SkM cells were transfected with 400 ng ABE mRNA and guide RNA at a 1:5 mRNA:gRNA molar ratio using Lipofectamine™ MessengerMAX™ (Invitrogen) following the manufacturer’s protocol. Cells were harvested three days post-transfection, and genomic DNA was extracted by adding 50 µL QuickExtract (LGC Biosearch Technologies) per well, mixing well, transferring to a PCR plate, and incubating at 65° for 30’, followed by 68° for 30’ and 98° for 10’. For SkM cells, samples were incubated at 65 °C for 6 minutes while shaking at 300 rpm prior to being transferred to PCR plates to ensure cells were fully lifted.

Target loci were PCR-amplified using Q5 High-Fidelity DNA Polymerase with gene-specific primers containing Illumina adapters, followed by a second PCR with barcoded primers for 10 cycles. Amplicons were sequenced on an Illumina MiSeq, and base editing frequencies were quantified using CRISPResso2.

### HEK293T cell fluorescence screen

Plasmids encoding the nominated ADA candidates were tested using a mammalian fluorescence-based screen. A pEditor plasmid encodes ABE-T2A-GFP in which an ABE and GFP reporter are coexpressed and separated by a T2A ribosomal skipping peptide, and a pReporter plasmid encodes BFP-T2A-target-mCherry that contains a target spacer sequence. In cells, an ABE expressed from a pEditor edits an in-frame stop codon to a sense codon on a pReporter, and mCherry expression acts as an indicator of ABE activity. HEK293T cells were transfected with Lipofectamine 2000 (120ng pEditor, 300ng pReporter) and MessengerMAX (100pmol synthetic gRNA). We used flow cytometry (Attune NxT) to measure the fraction of dually transfected cells, which is the sub-population positive for both GFP and BFP, that also expressed mCherry. This fraction of mCherry-positive cells in the dually transfected sub-population is increased for ABE variants that have improved activity. Synthetic gRNAs containing phosphorothioate and 2’-O-methyl chemical modifications were synthesized by IDT **(Table S3).**

## Author contributions

K.L.R. wrote the manuscript with input from all authors. K.L.R., J.M.L.H, I.N, J.W.T., N.C.T, S.J., C.J.C., J-L L., B.F., L.G-O., J.R. designed and performed the experiments, and analyzed the data that enabled the discovery and mutagenesis of improved variants of deaminase and nuclease of the base editors. S.T. purified protein for structural analysis. R.F.O, J.T.W., and I.K. determined the protein structures. K.S. and B.J. designed, performed, and analyzed the DMD editing experiments. L.M.A., A.R.B, C.T.B, D.S.A.G., K.G.H., P.S., B.C.T, D.W.T, and C.N.B. guided the research.

## SUPPLEMENTARY INFORMATION

### Supplementary Note 1: Development of MG102-71 via Ancestral Reconstruction

Type II-D nucleases are compact and compatible with AAV delivery but are less prevalent than Type II-A nucleases and often display limited activity in mammalian cells in their wild-type form. We previously applied ancestral sequence reconstruction (ASR) to enhance activity and diversify MG34 Cas9d nucleases. Given the improved performance of reconstructed MG34 variants **[1]**, we applied a similar strategy to the MG102 Cas9d clade.

MG102-2, MG102-39, MG102-14, and MG102-35 were previously identified and characterized, with MG102-2 exhibiting the highest activity in vitro and in mammalian cells **[2]**. These extant nucleases were used to infer putative ancestral sequences using maximum-likelihood phylogenetic methods.

A total of 441 MG102 homologs were aligned using MAFFT (G-INS-i) **[3]**, and a phylogenetic tree was constructed **(Fig. S1A).** The tree was rooted using SpCas9 and SaCas9 as outgroups. Ancestral sequences were inferred using PAML v4.8 **[4]**. Insertions and deletions were manually curated for each reconstructed node.

To generate additional ancestral intermediates, amino acid substitutions were introduced between reconstructed nodes using a weighted probabilistic approach. Global weights were assigned to each ancestor **(Table S1)**, and posterior probabilities for all twenty amino acids at each position were used to guide residue selection during recombination **(Fig. S1A).** Eighteen ancestral variants were generated within the MG102 clade; five exhibited detectable in vitro nuclease activity **(Fig. S1A, C).**

Variants were evaluated in K562 cells using the MG102-2 sgRNA scaffold across eight genomic loci adjacent to NRC PAMs, where MG102-2 displayed modest activity. MG102-71 showed activity at all eight loci and was functional with both MG102-2 and MG102-39 sgRNA scaffolds. MG102-68 exhibited comparable activity.

PAM profiling of MG102-71 by NGS of a randomized PAM library confirmed retention of an NRC PAM preference despite only sharing 73% sequence identity within the PAM-interacting domain relative to MG102-2.

### Supplementary Note 2: Development and engineering of an MG3-6–based ABE platform

We mined large-scale metagenomic datasets to identify novel adenosine deaminases (ADAs). From this search, we identified a clade of candidate enzymes closely related to EcTadA, which we designated as the MG68 family. Several MG68 candidates were evaluated as N-terminal fusion constructs with SpCas9 nickases, using EcTadA-based ABE as a positive control. Among the tested variants, three candidates, MG68-3, MG68-4, and MG68-5, demonstrated detectable deamination activity in our screening assay **(Fig. S2)**. Of these, MG68-4 exhibited the most favorable activity profile and was therefore selected for further engineering and optimization.

Previously, we reported the discovery, characterization, and engineering of the Type II-A nuclease MG3-6_3-8, which exhibits high indel activity coupled with strong target specificity **[5]**. Given its strong activity and genome-wide targeting capacity enabled by PID chimeragenesis **[6],** we aimed to repurpose MG3-6_3-8 as a modular ABE platform and used it as the scaffold for systematic engineering and optimization of the MG68-4.

To construct ABEs based on MG68-4, we fused two copies of the MG68-4 adenosine deaminase, connected by a 65-amino-acid linker. We inserted this cassette into MG3-6_3-8 nickase (D13A) to generate a panel of inlaid ABE architectures. Specifically, the tandem MG68-4 cassette was inserted at 30 distinct positions across the nuclease scaffold, with particular emphasis on the REC, HNH, and RuvC domains and using varying linker lengths **(Fig. S3A).** A complete summary of all fusion constructs is provided in Table S1. Insertion sites within the WED and PID domains were intentionally avoided, as these regions serve as chimeric boundaries for PID swapping with homologous MG3 nucleases, and we wanted to preserve the chimeragenesis capability of the platform **[5, 6].**

All ABE architectures were evaluated in HEK293T cells using a spacer targeting the TRAC locus containing multiple adenines. This locus was selected based on the previously observed high editing efficiency of MG3-6_3-8 **[5]**, thereby enabling concurrent evaluation of overall editing activity and editing window properties. Terminal fusions of the double-copy MG68-4 cassette exhibited minimal base editing activity, with C-terminal fusions outperforming

N-terminal fusions. In contrast, several inlaid architectures demonstrated robust activity, particularly insertions within the REC domain (site I297) and multiple positions within the RuvC-III domain **(Table S2, Fig. S3B).** Among these RuvC architectures, the inlay at position L792 was selected as the lead architecture and is hereafter referred to as ABE07.

We next leveraged ABE07 as a chassis to further engineer the MG68-4 deaminase for improved deamination performance **(Table S3).** Editing activity was assessed in Hepa1-6 cells across a panel of 15 guides to enable robust benchmarking of variant performance **(Table S4).**

Because our ultimate objective was to obtain a single optimized deaminase variant compatible with both MG34-29 and MG102-71, we first asked whether one or both deaminase copies were responsible for base editing activity in ABE07. *Ec*TadA is known to form an obligate dimer, and we hypothesized that the novel MG68-4 shared this property. Accordingly, these dimeric architectures were designed to emulate the native oligomeric configuration of the deaminase domain. To minimize editor size and dissect the functional contribution of each deaminase subunit, we generated a panel of monomeric (ABE01, ABE02, ABE55), homodimeric (ABE51, ABE54, ABE58), and heterodimeric (ABE51, ABE52, ABE57) ABE variants **(Table S3)**.

To probe subunit-specific activity in more detail, we selectively deactivated(via an E60A mutation) or completely removed either the N-terminal or C-terminal deaminase catalytic domain (ABE51, ABE52, ABE54, and ABE57). Comparative analysis revealed that base editing activity was dependent exclusively on the C-terminal deaminase subunit. Variants with an inactivated C-terminal domain (ABE52) behaved similarly to fully inactive controls (ABE51), indicating that the N-terminal deaminase is not positioned to access the exposed single-stranded DNA substrate. Conversely, variants with an inactivated N-terminal subunit (ABE54 and ABE57) retained substantial editing activity and frequently outperformed monomeric and homodimeric constructs **(Fig. S3B, S3C)**.

Collectively, these results demonstrate that the C-terminal deaminase domain is solely responsible for genomic DNA editing, while the N-terminal domain may provide structural or stabilizing support. This observation suggested that the functional contribution of the heterodimeric configuration could potentially be compensated for through targeted mutagenesis of a single deaminase domain.

We next sought to enhance the activity of the heterodimeric ABE15 variant by introducing additional mutations into the C-terminal deaminase subunit. Building upon a prior combinatorial mutagenesis screening, we generated variants ABE32-ABE41 **(Table S3)** and screened them on the 15 guide set **(Table S4)**. Among these, incorporation of the L85F mutation alongside the existing D109N, T112R, and A155R mutations (ABE33) produced the largest improvement in editing performance. ABE33 exhibited a mean A→G editing efficiency of 32.97%, representing a ∼1.5-fold increase over ABE15 (22.8% mean editing; Fig. S3A). Notably, ABE33 also enabled near-saturating editing across multiple adenines within individual protospacers, exhibiting both a higher maximum editing efficiency and reduced deviation compared to other variants **(Fig. S4A, B).**

In contrast, incorporation of A143W (ABE34), A155E (ABE35), E10Y (ABE37), or A126D (ABE40) reduced A→G editing efficiency **(Fig. S4A).** These results identify L85F as a key mutation that enhances MG68-4 deaminase performance within the MG3-6 ABE chassis.

While analyzing ABE15 activity across the 15 guide panel, we observed that the MG68-4 deaminase exhibits an intrinsic preference for a -NAC- motif, reminiscent of the -UACG- motif recognized by the tRNA adenosine deaminases **(Fig. S4C) [5–9]**. Although the tested spacers contained numerous adenines flanked by guanines at both the −1 and +1 positions **(Fig. S4C)**, ABE15 preferentially edited adenines with a +1 cytosine compared to those with a +1 guanine **(Fig. S4B)**. To relax this sequence context bias and broaden the editable sequence space, we designed a series of proline substitutions within the terminal α-helix of the deaminase domain **(Table S3)**. We hypothesized that these substitutions would increase local conformational flexibility and expand the active site to better accommodate purine bases. Among the variants tested, simultaneous incorporation of R153P and R154P mutations into the ABE15 background (ABE23) resulted in increased mean and maximum A→G editing efficiencies, without a concomitant increase in cytosine deamination or indel formation. ABE23 achieved a mean A→G editing efficiency of 30.8%, compared to 21.4% for ABE15, and a higher mean maximum editing efficiency (82.1% versus 73.5%) across the 15 guides tested **(Fig. S4D, E).** Importantly, the enhanced activity of ABE23 was accompanied by a relaxation of the stringent +1 cytosine sequence preference observed in ABE15 **(Fig. S4F)**, indicating successful broadening of the deaminase editing context.

### Supplementary Note 3: Fluorescent Reporter Assay for High-Throughput Base Editor Screening

Since most deaminase engineering was performed in Hepa1-6 cells and we envision using compact base editors in mammalian systems, we developed a fluorescence-based reporter assay in HEK293T cells to enable high-throughput evaluation of engineered MG68-4 deaminase variants in a human cellular context **(Fig. S5A)**. In this system, cells were co-transfected with an ABE expression construct (ABE–T2A–GFP), a fluorescent reporter plasmid in which a premature stop codon disrupts mCherry expression, and a targeting sgRNA. Successful A→G editing of the stop codon restores mCherry fluorescence, allowing direct quantification of editing efficiency by flow cytometry within the doubly transfected (GFP⁺/BFP⁺) cell population **(Fig. S5A).**

Given that multiple deaminase variants **(Table S3)** exhibited comparable activity across different adenines and sequence contexts in prior screens, we selected a focused set of recurrently beneficial residues identified in mammalian assays, namely V83S, L85F, D109N, T112R, R153P, R154P, and A155R, for combinatorial optimization within the MG3-6_3-8 ABE framework **(Fig. S5B, C)**. These residues were systematically recombined and screened using the fluorescence reporter assay to identify high-performance MG68-4 variants under this orthogonal selection modality. This screen identified two top-performing ABE variants, ABE66 and ABE70, which consistently exhibited the highest fractions of mCherry-positive cells among the tested constructs.

The engineered deaminase domains from these variants (monomeric MG68-4(V83S, D109N, T112R, R154P) in ABE66 and monomeric MG68-4(V83S, D109N, T112R, R153P, R154P) in ABE70) were subsequently nominated for downstream protein engineering of SMART ABEs in HEK293T cells **(Fig. S5D, E)**.

## Supplementary Tables

**Table S1:** Ancestral Sequence Reconstruction (ASR) Posterior Probabilities and Sequence Proximity to MG102-2.

| MG code | Mean support | %ID to MG102-2 |
| --- | --- | --- |
| **MG102-68** | 0.91 | 79.25% |
| **MG102-71** | 0.91 | 76.37% |
| **MG102-77** | 0.89 | 54.32% |
| **MG102-78** | 0.87 | 58.12% |
| **MG102-79** | 0.88 | 69.41% |

**Table S2:**
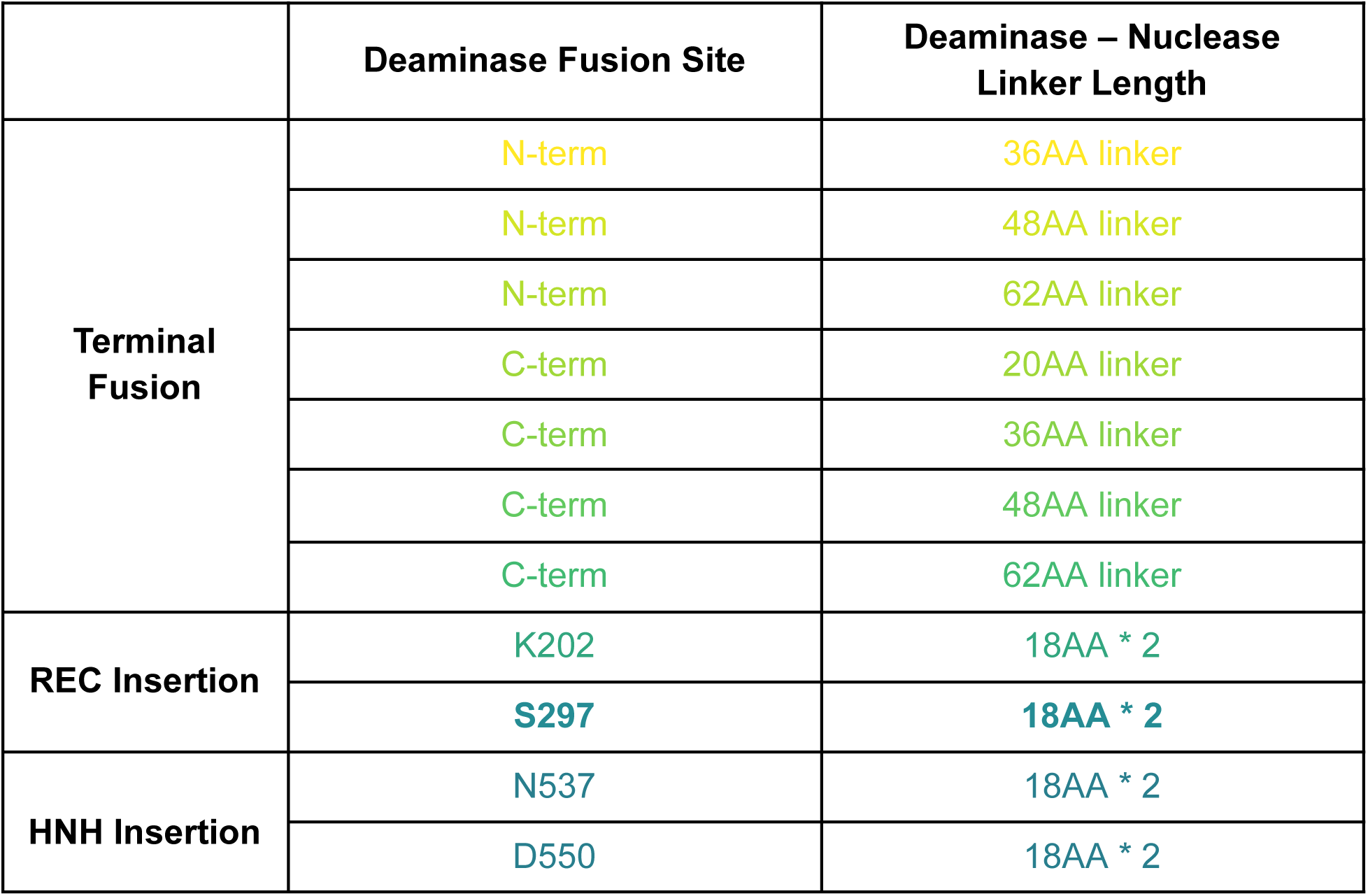
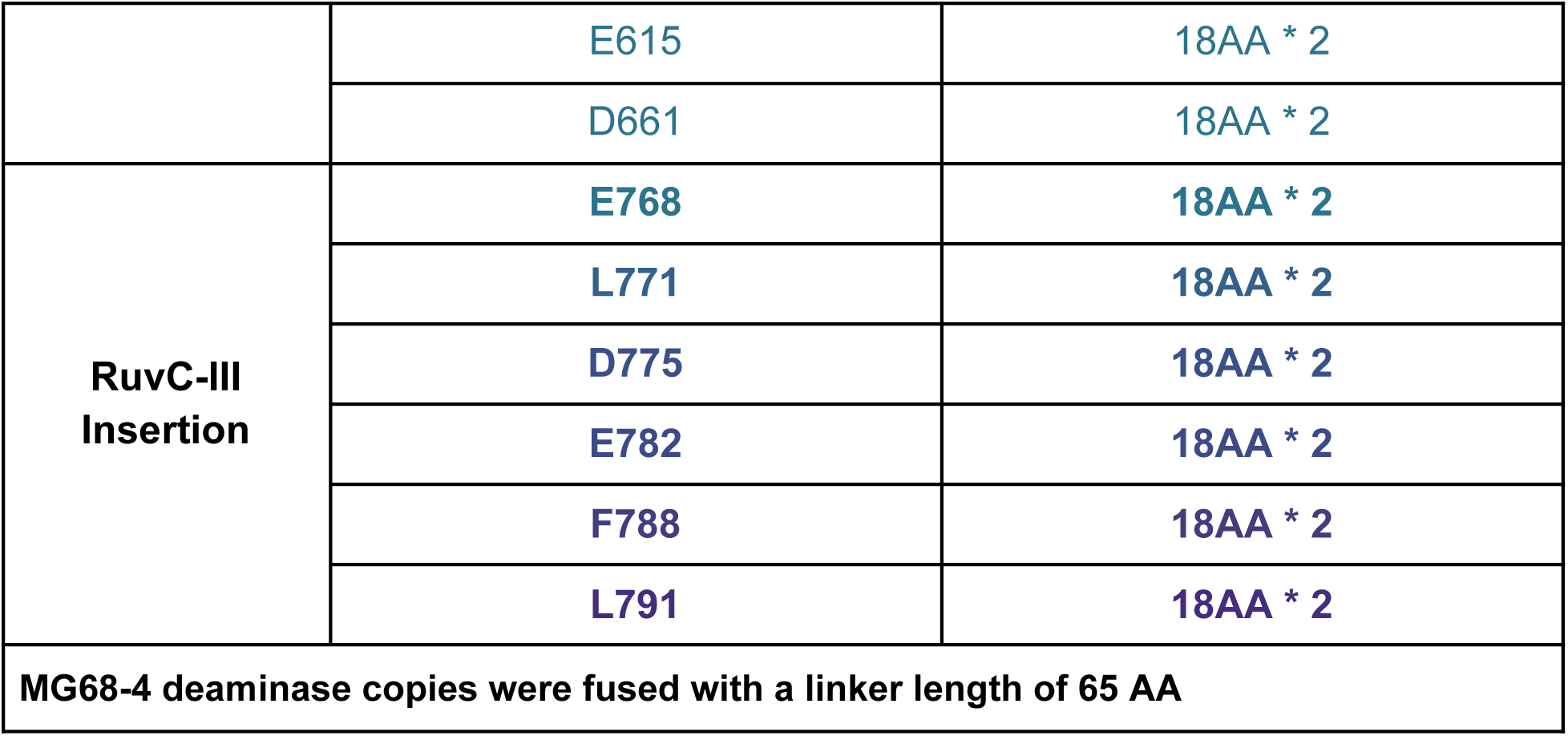
MG3-6_3-8 and MG68-4 fusion architectures tested in this study.

**Table S3:** Genotypes of MG3-6_3-8 ABEs described in this study. The most significant ABE variants designed are highlighted in bold.

| **ABE variant** | **Architecture** | **Deaminase Mutations (N-term + C-term)** |
| --- | --- | --- |
| ABE01 | Monomeric | MG68-4 (D109N) - inlaid in RuvC domain |
| ABE02 | Monomeric | MG68-4 (D109N) - inlaid in Rec domain |
| ABE07 | Homodimeric | MG68-4 (D109N)<br>+ MG68-4 (D109N) |
| ABE51 | Heterodimeric | MG68-4 (D109N)<br>+ dead MG68-4 |
| ABE52 | Heterodimeric | dead MG68-4<br>+ MG68-4 (D109N) |
| ABE54 | Homodimeric | dead MG68-4<br>+ dead MG68-4 |
| ABE55 | Monomeric | MG68-4 (D109N+T112R+A155R) |
| ABE57 | Heterodimeric | dead MG68-4<br>+ MG68-4 (D109N+T112R+A155R) |
| ABE58 | Homodimeric | MG68-4 (D109N+T112R+A155R)<br>+ MG68-4 (D109N+T112R+A155R) |
| **ABE15** | **Heterodimeric** | **wt MG68-4** |
|  |  | **+ MG68-4 (D109N+T112R+A155R)** |
| **ABE32** | **Heterodimeric** | **wt MG68-4**<br>**+ MG68-4 (V83S+D109N+T112R+A155R)** |
| **ABE33** | **Heterodimeric** | **wt MG68-4**<br>**+ MG68-4 (L85F+D109N+T112R+A155R)** |
| ABE34 | Heterodimeric | wt MG68-4<br>+ MG68-4 (A143W+D109N+T112R+A155R) |
| ABE35 | Heterodimeric | wt MG68-4<br>+ MG68-4 (A155E+D109N+T112R) |
| ABE36 | Heterodimeric | wt MG68-4<br>+ MG68-4 (A160T+D109N+T112R+A155R) |
| ABE37 | Heterodimeric | wt MG68-4<br>+ MG68-4 (E10Y+D109N+T112R+A155R) |
| ABE38 | Heterodimeric | wt MG68-4<br>+ MG68-4 (D162Q+D109N+T112R+A155R) |
| ABE39 | Heterodimeric | wt MG68-4<br>+ MG68-4 (S147F+D109N+T112R+A155R) |
| ABE40 | Heterodimeric | wt MG68-4<br>+ MG68-4 (A126D+D109N+T112R+A155R) |
| ABE41 | Heterodimeric | wt MG68-4<br>+ MG68-4 (F150C+D109N+T112R+A155R) |
| **ABE23** | **Heterodimeric** | **wt MG68-4**<br>**+MG68-4(D109N+T112R+R153P+R154P+A155R)** |
| ABE24 | Heterodimeric | wt MG68-4<br>+ MG68-4 (D109N+T112R+R153P+A155R) |
| ABE25 | Heterodimeric | wt MG68-4<br>+ MG68-4 (D109N+T112R+R154P+A155R) |
| ABE26 | Heterodimeric | wt MG68-4<br>+ MG68-4 (D109N+T112R+delR154+A155R) |
| ABE60 | Monomeric | MG68-4 (V83S+D109N+R154P) |
| ABE61 | Heterodimeric | wt MG68-4<br>+ MG68-4 (V83S+D109N+R154P) |
| ABE62 | Monomeric | wt MG68-4<br>+ MG68-4 (V83S+D109N+R153P) |
| ABE63 | Heterodimeric | MG68-4 (V83S+D109N+R153P) |
| ABE64 | Monomeric | MG68-4 (V83S+D109N+R153P+R154P+A155R) |
| ABE65 | Heterodimeric | wt MG68-4<br>+ MG68-4 (V83S+D109N+R153P+R154P+A155R) |
| **ABE66** | **Monomeric** | **MG68-4 (V83S+D109N+T112R+R154P)** |
| ABE67 | Heterodimeric | wt MG68-4<br>+ MG68-4 (V83S+D109N+T112R+R154P) |
| ABE68 | Monomeric | MG68-4 (V83S+D109N+T112R+R153P+A155R) |
| ABE69 | Heterodimeric | wt MG68-4<br>+ MG68-4 (V83S+D109N+T112R+R153P+A155R) |
| **ABE70** | **Monomeric** | **MG68-4 (V83S+D109N+T112R+R153P+R154P)** |
| ABE71 | Heterodimeric | wt MG68-4<br>+ MG68-4 (V83S+D109N+T112R+R153P+R154P) |

**Table S4.**
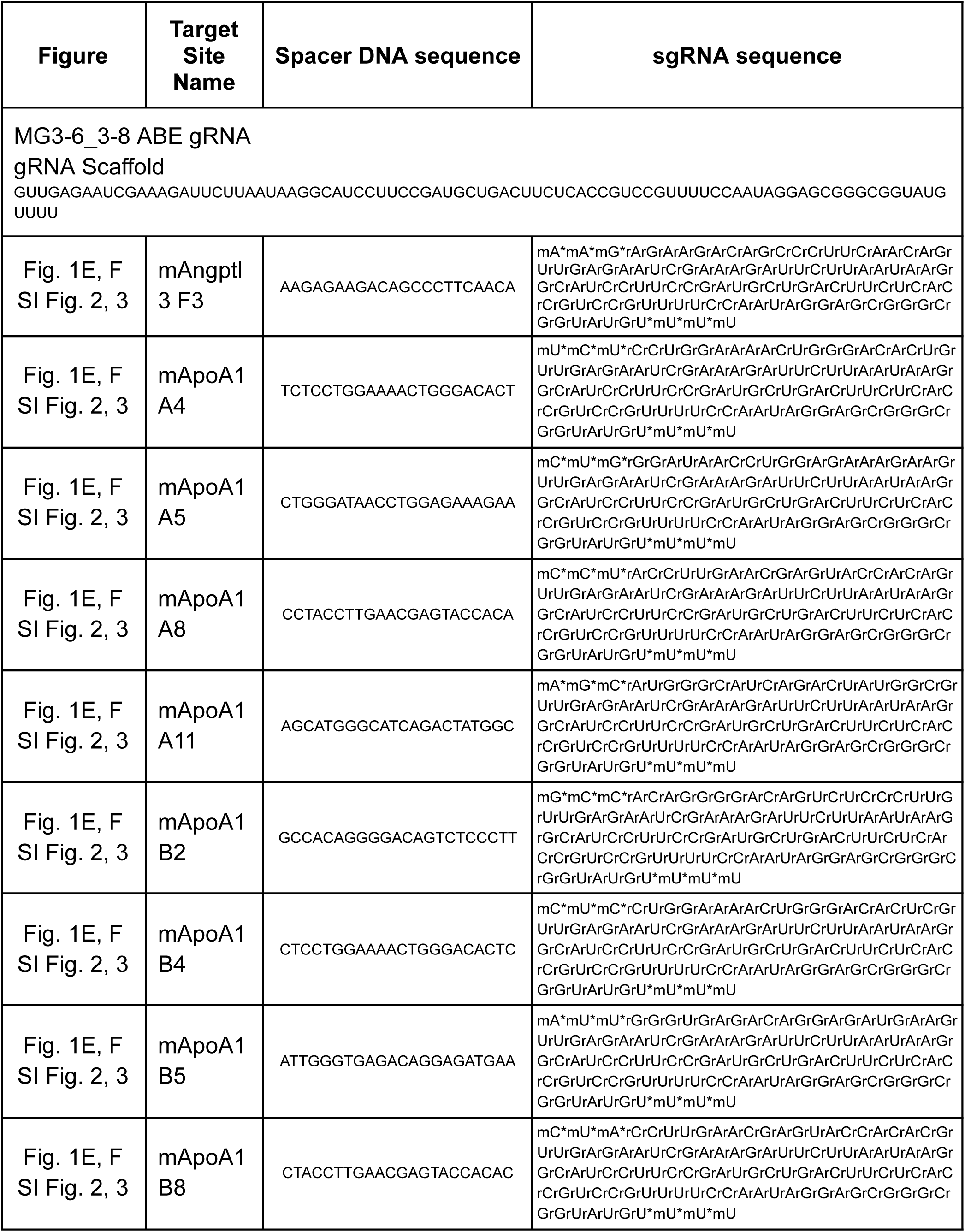
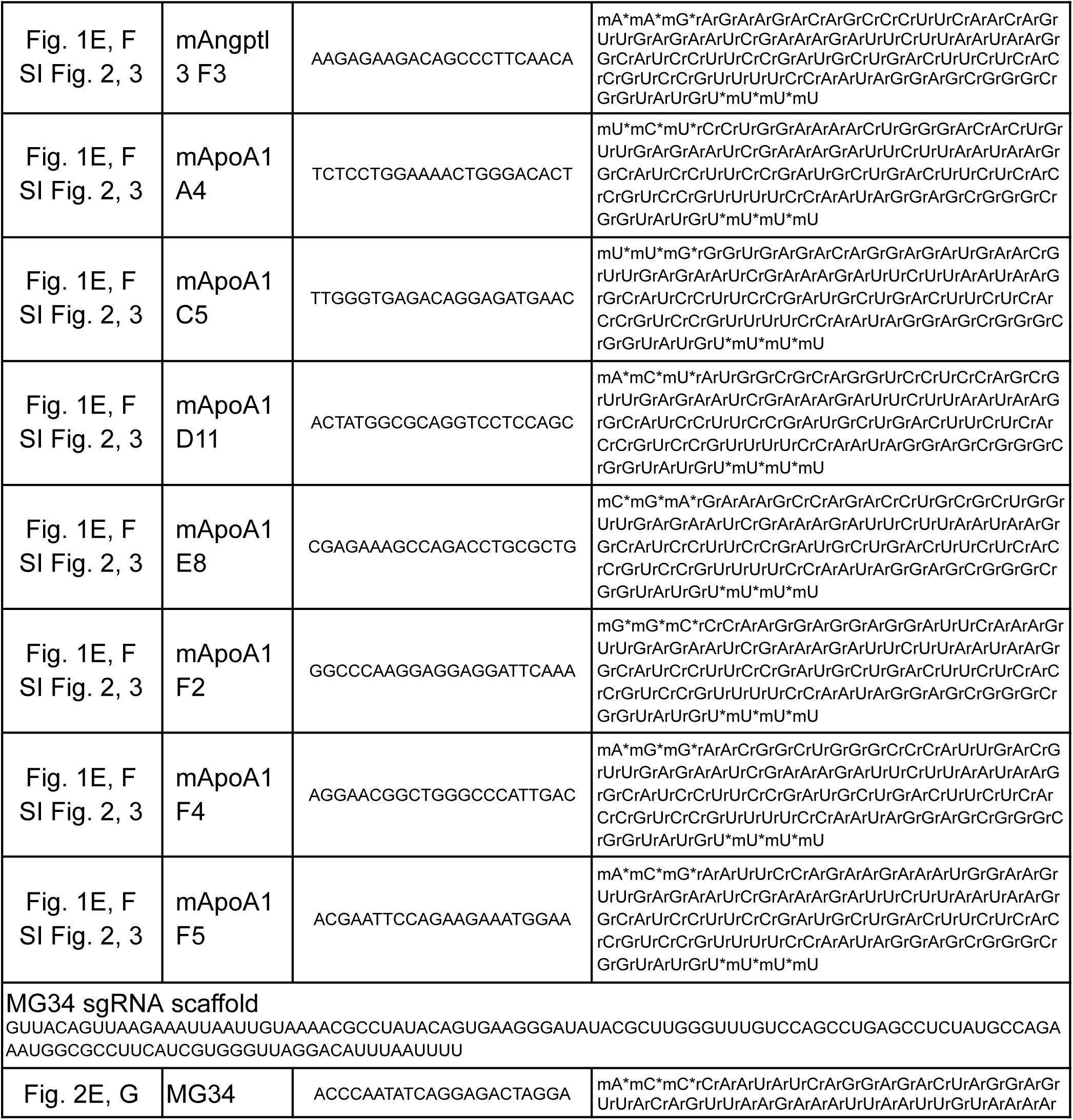
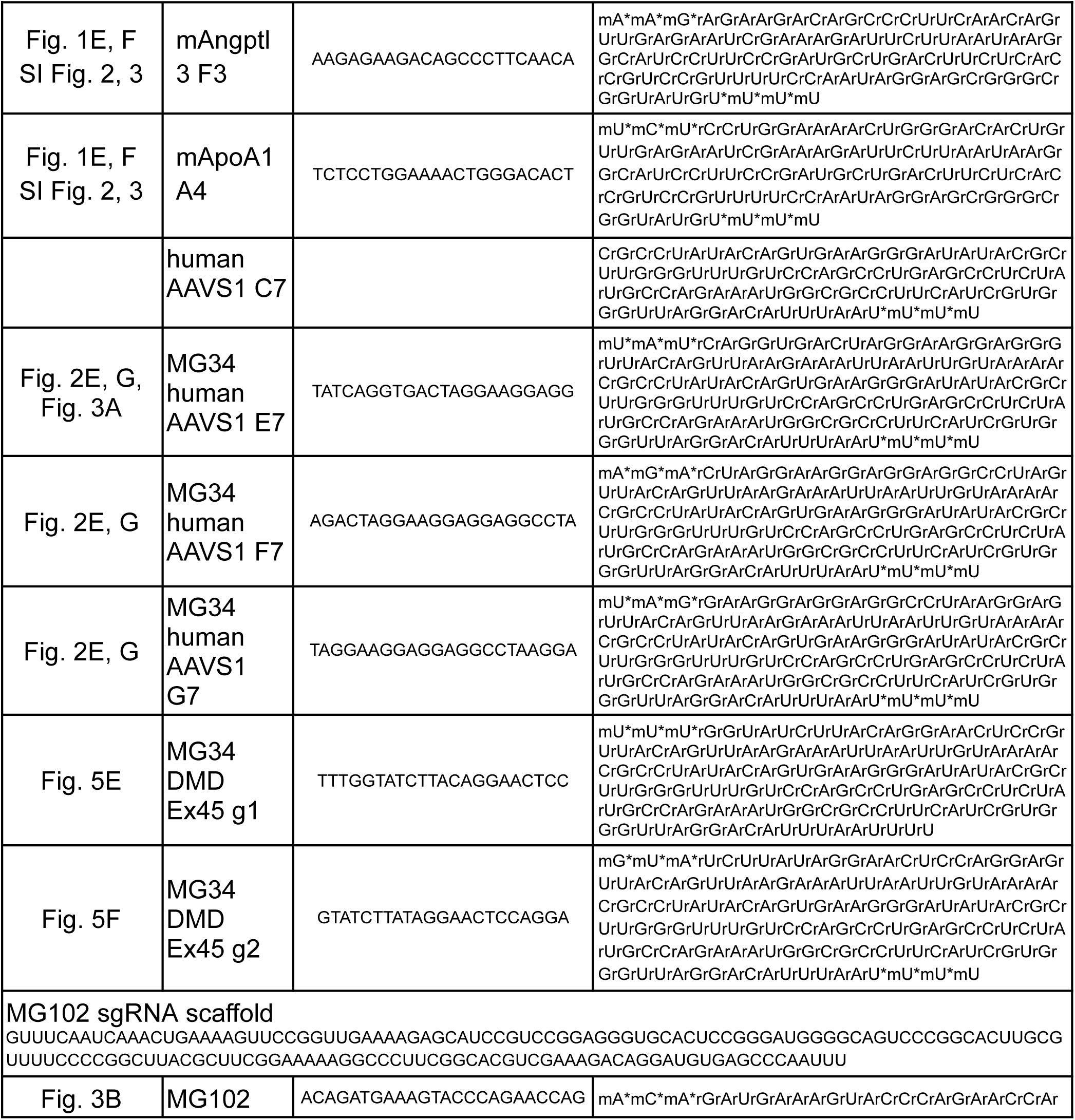
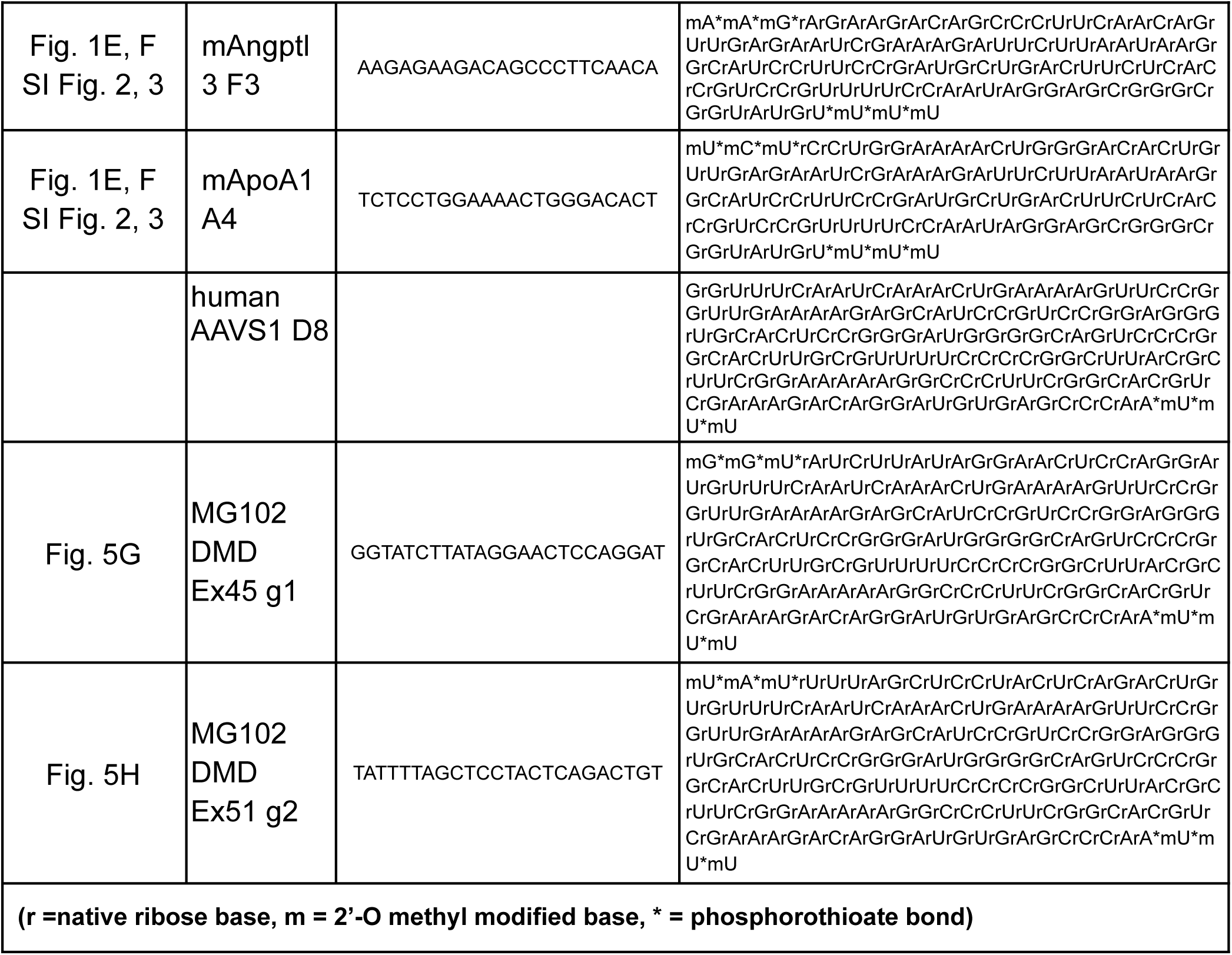
sgRNA sequences used in this manuscript.

## Supplementary Figures

**Fig. S1:**
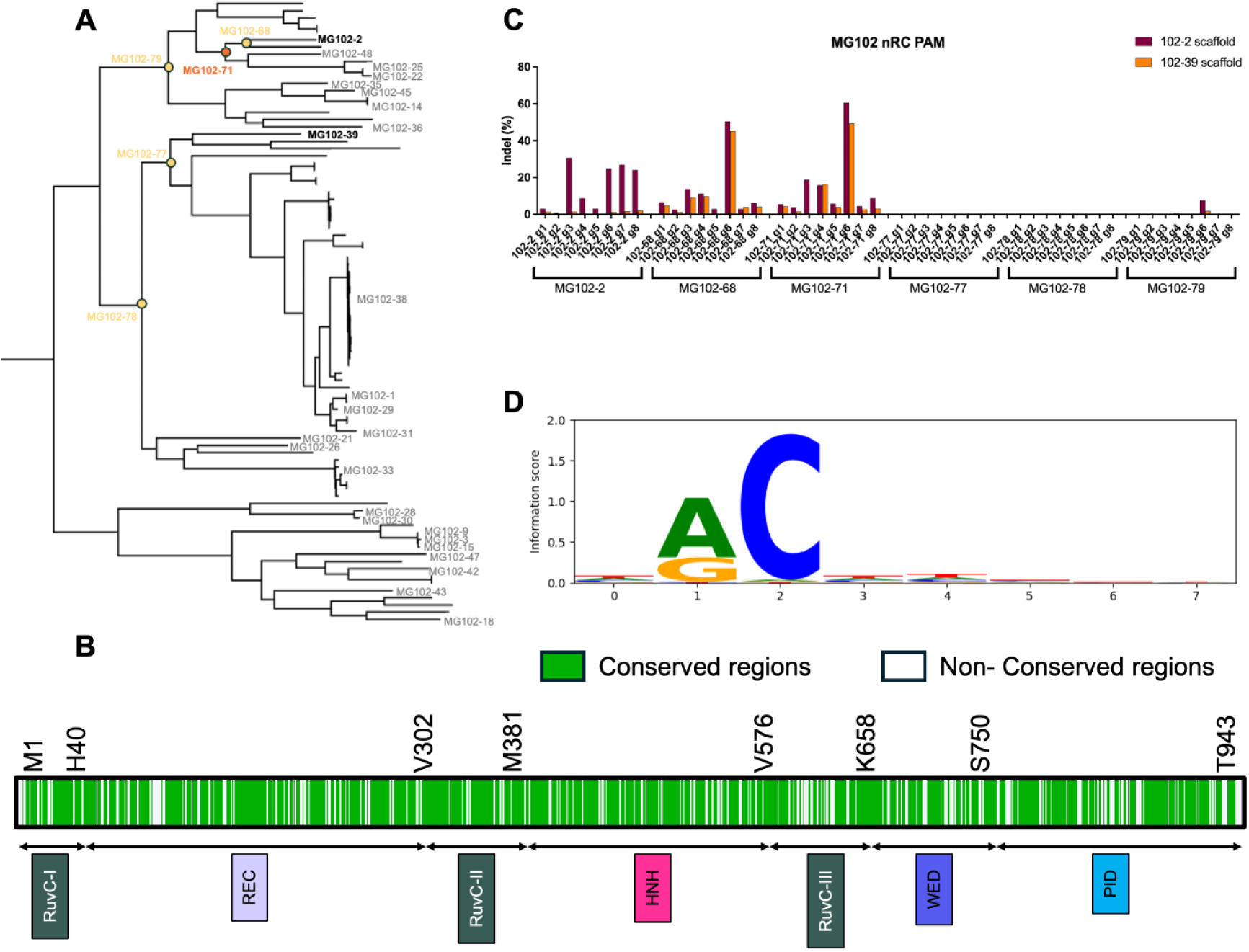
Development of MG102-71 via Ancestral Reconstruction. **(A)** Maximum-likelihood phylogenetic tree of MG102 Cas9d homologs generated using MAFFT-linsi multiple sequence alignment and RAxML. The reconstructed ancestral protein is indicated by the circle. Scale bar denotes substitutions per site. **(B)** Pairwise protein sequence alignment of MG102-2 and MG102-71. Conserved residues are shown in green and non-conserved residues are shown in white. **(C)** PAM SeqLogo of MG102-71 based on NGS of cleaved plasmids from a plasmid PAM library tested with MG102-2 guide scaffold. **(D)** Indel editing efficiencies of ancestral MG102 variants in K562 cells using the wild-type MG102 sgRNA scaffold across 8 spacers targeting AAVS1 loci adjacent to NRC PAMs. Values represent the mean of two biological replicates.

**Fig. S2:**
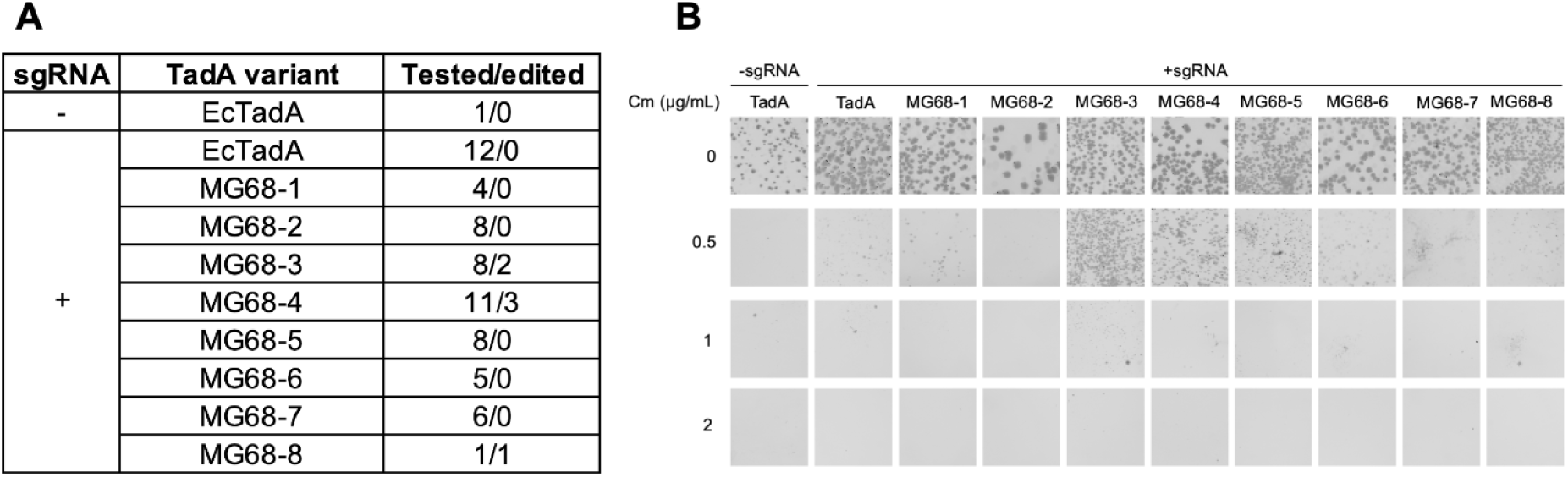
Investigation and characterization of MG ADA activity through the positive selection. **(A)** Editing efficiencies of MG ADA candidates. MG68-3, MG68-4, and MG68-5 showed base edits of adenine. Colonies were picked from the plates with greater than or equal to 0.5 μg/mL Cm. **(B)** ABEs are composed of N-terminal ADA variants and C-terminal SpCas9 (D10A) nickase. Eight MG68 ADAs were tested against 0 to 2 μg/mL of chloramphenicol. For simplicity, only the identities of deaminases are shown.

**Fig. S3:**
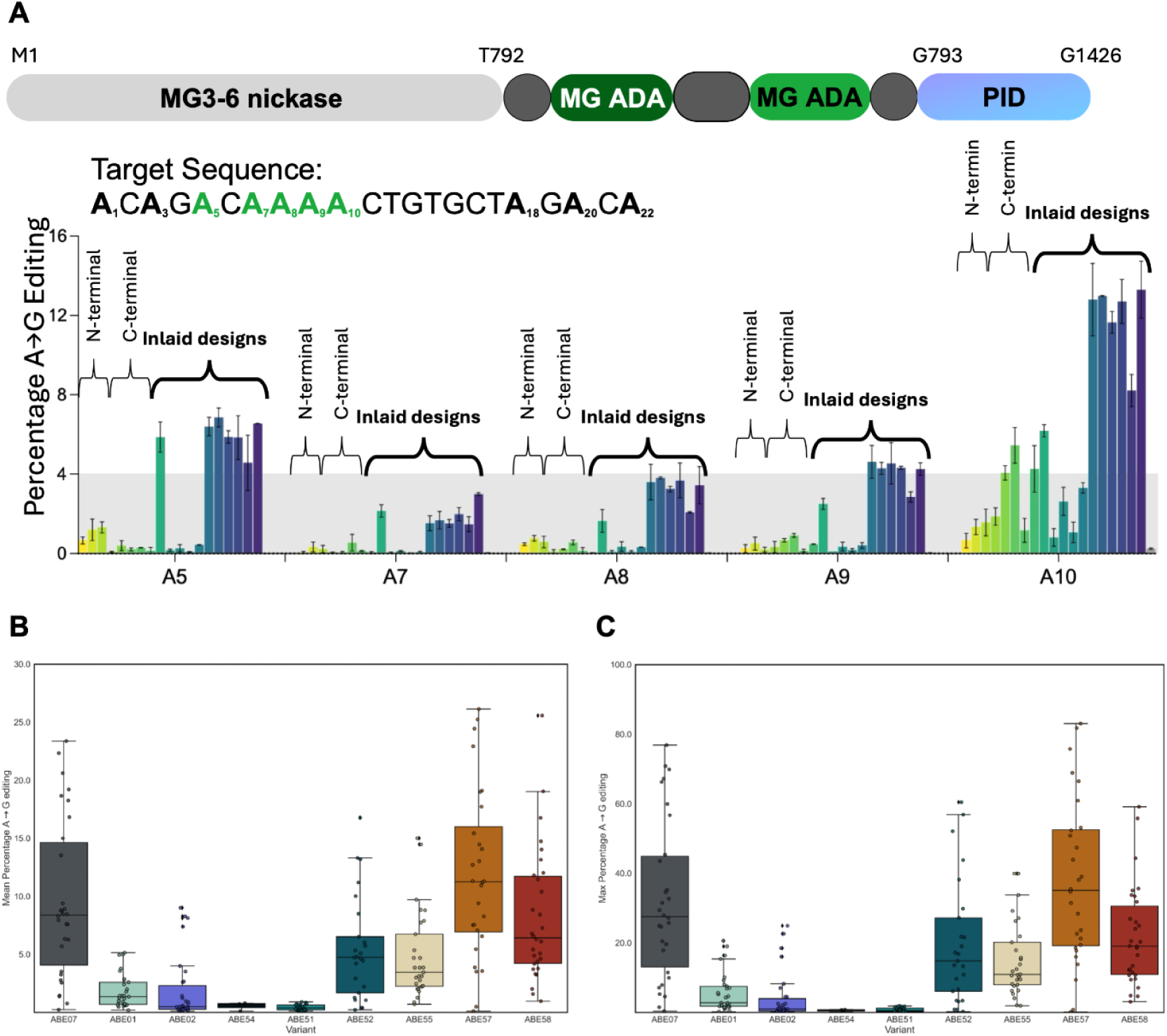
Optimization of MG3-6_3-8 and MG68-4 ABE architecture. **(A)** Schematic of MG3-6_3-8 ABE architectures and comparison of terminal versus inlaid MG68-4 deaminase fusion strategies. Two copies of the MG68-4 adenosine deaminase were fused to the MG3-6_3-8 nickase either at the N- or C-terminus or inlaid at multiple internal positions. The residue numbering is relative to the MG3-6_3-8 nuclease. Bar plots show A→G editing efficiencies at individual adenines (A5–A10) within a representative target sequence. Boxplots depicting **(B)** Mean observed editing and **(C)** Max observed editing by individual ABE variants to oligomeric variants of 15 distinct gRNA targeting 2 different genes in Hepa1-6 cells. Editing data corresponding to n= 2 biologically independent replicates for each target guide. Individual data points represent each unique target’s locus.

**Fig S4:**
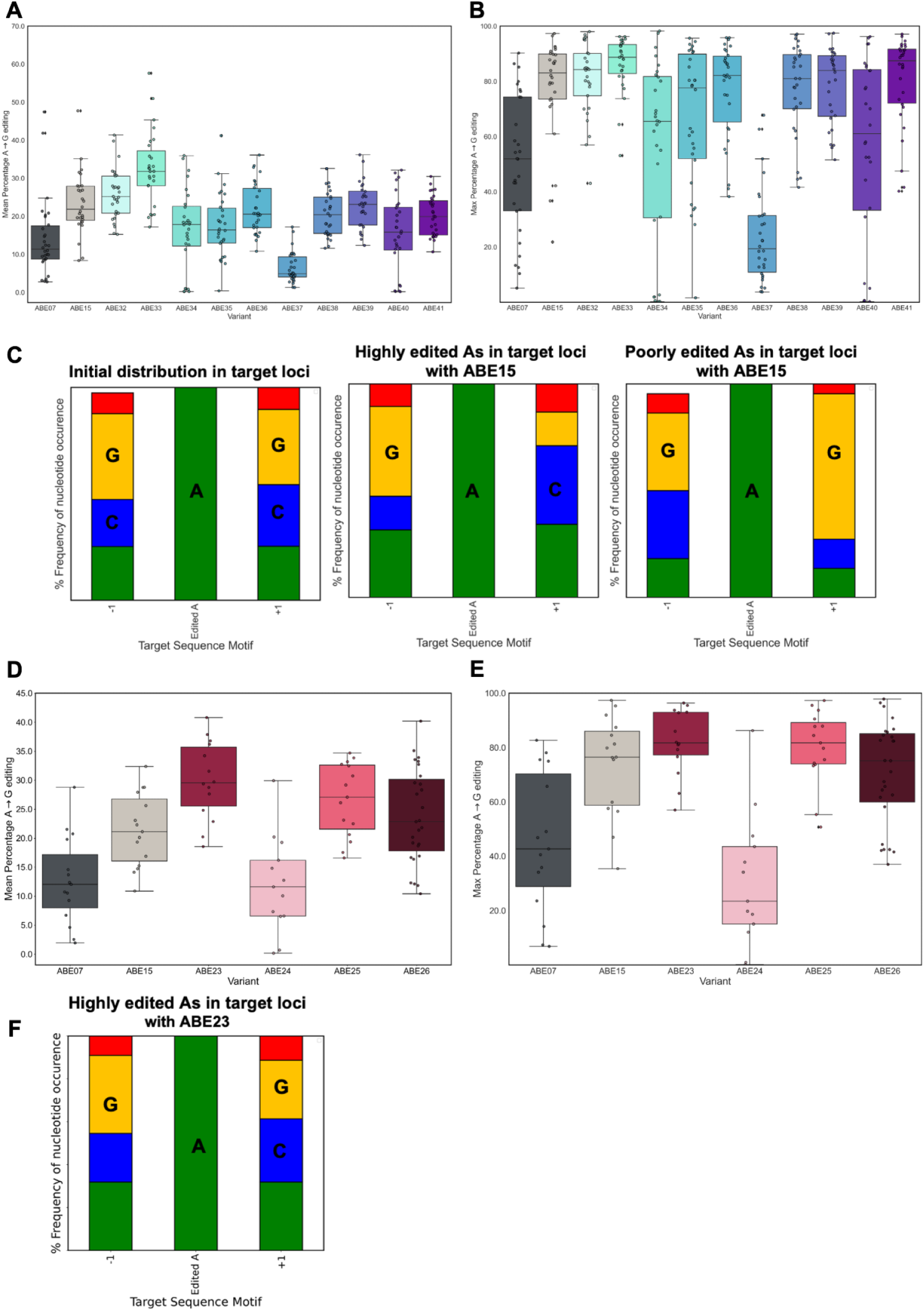
Engineering of the MG68-4 deaminase within the MG3-6_3-8 ABE platform in Hepa1-6 cells. Boxplots showing **(A)** Mean and **(B)** Max A→G editing efficiencies for single point mutants of MG68-4. **(C)** Sequence context preferences of edited adenines. Frequency distributions of nucleotides flanking the edited adenine (−1 and +1 positions) are shown for all editable adenines across the 15 gRNAs tested (left), highly edited adenines (>30% editing) by ABE15 (middle), and poorly edited adenines (<30% editing) by ABE15 (right). Boxplots showing **(D)** Mean and **(E)** Max A→G editing efficiencies of engineered proline-substituted MG68-4 variants evaluated across the same 15-gRNA panel. **(F)** Frequency distribution of nucleotides flanking highly edited adenines (>30% editing) by ABE23, illustrating relaxation of sequence context preference relative to ABE15.

**Fig S5:**
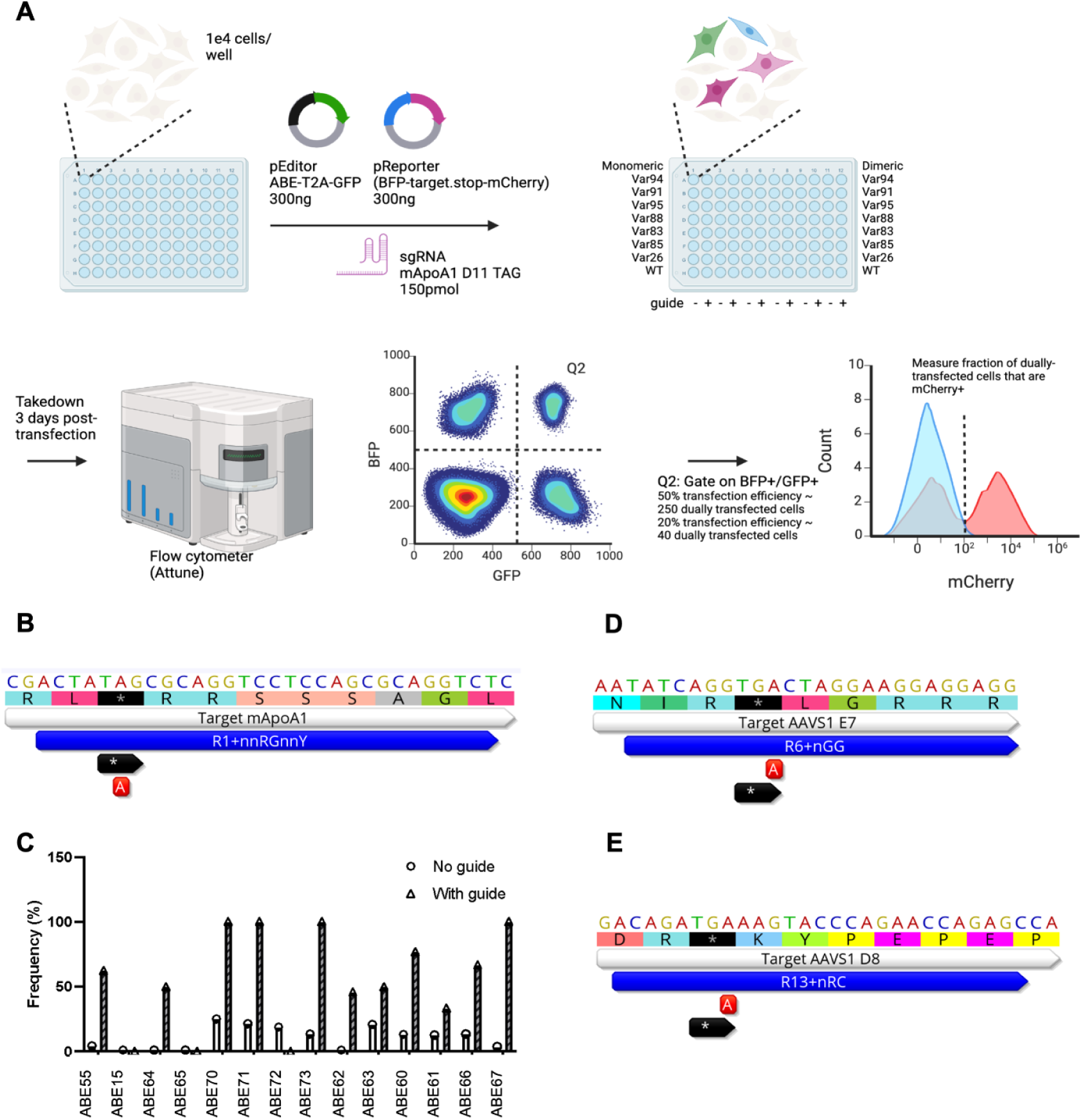
Molecular basis of fluorescence-based HEK293T screening of MG68-4 deaminase variants. **(A)** Schematic of the mammalian fluorescence-based screening assay. HEK293T cells were co-transfected with an ABE expression plasmid (pEditor, ABE-T2A-GFP), a reporter plasmid containing a stop codon–interrupted mCherry cassette (pReporter, BFP-target-stop-mCherry), and a targeting sgRNA. Cells were analyzed by flow cytometry three days post-transfection. Dual GFP/BFP gating was used to identify successfully transfected cells, and ABE activity was quantified by the fraction of mCherry-positive cells. ABE converts the target adenine in an in-frame stop codon of the target **(B)** mApoA1 in Hepa1-6 cells or **(D, E)** AAVS1 in HEK293T cells to a TGG sense codon, which enables expression of the mCherry protein. **(C)** Frequency of recovered MG3-6_3-8 ABE variants from the selection scheme with and without guide RNA, demonstrating guide-dependent enrichment of high-activity deaminase variants.

**Fig. S6:**
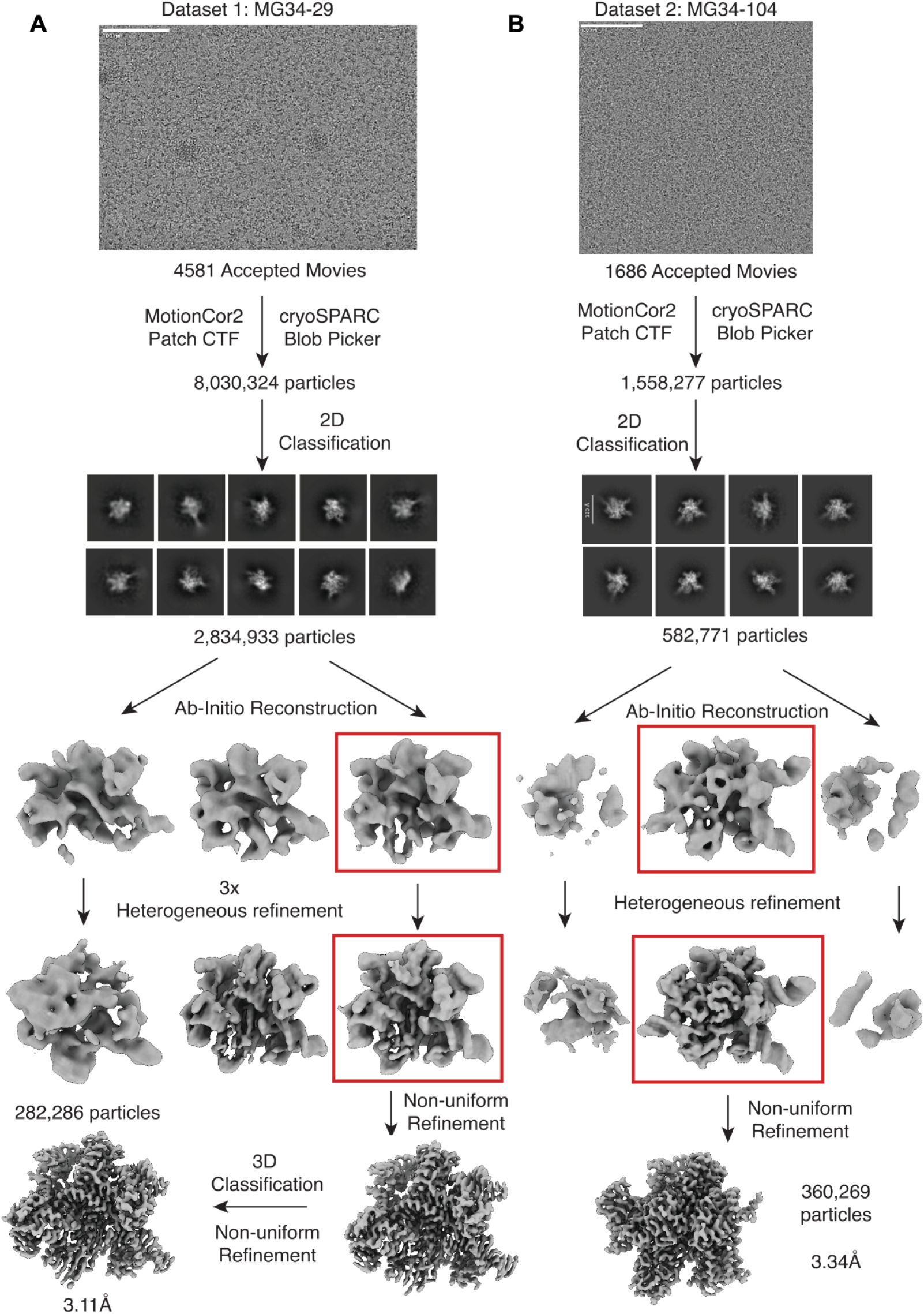
Cryo-EM dataset processing pipeline. Processing pipeline for (A) MG34-29 and (B) MG34-104 ternary complexes.

**Fig. S7:**
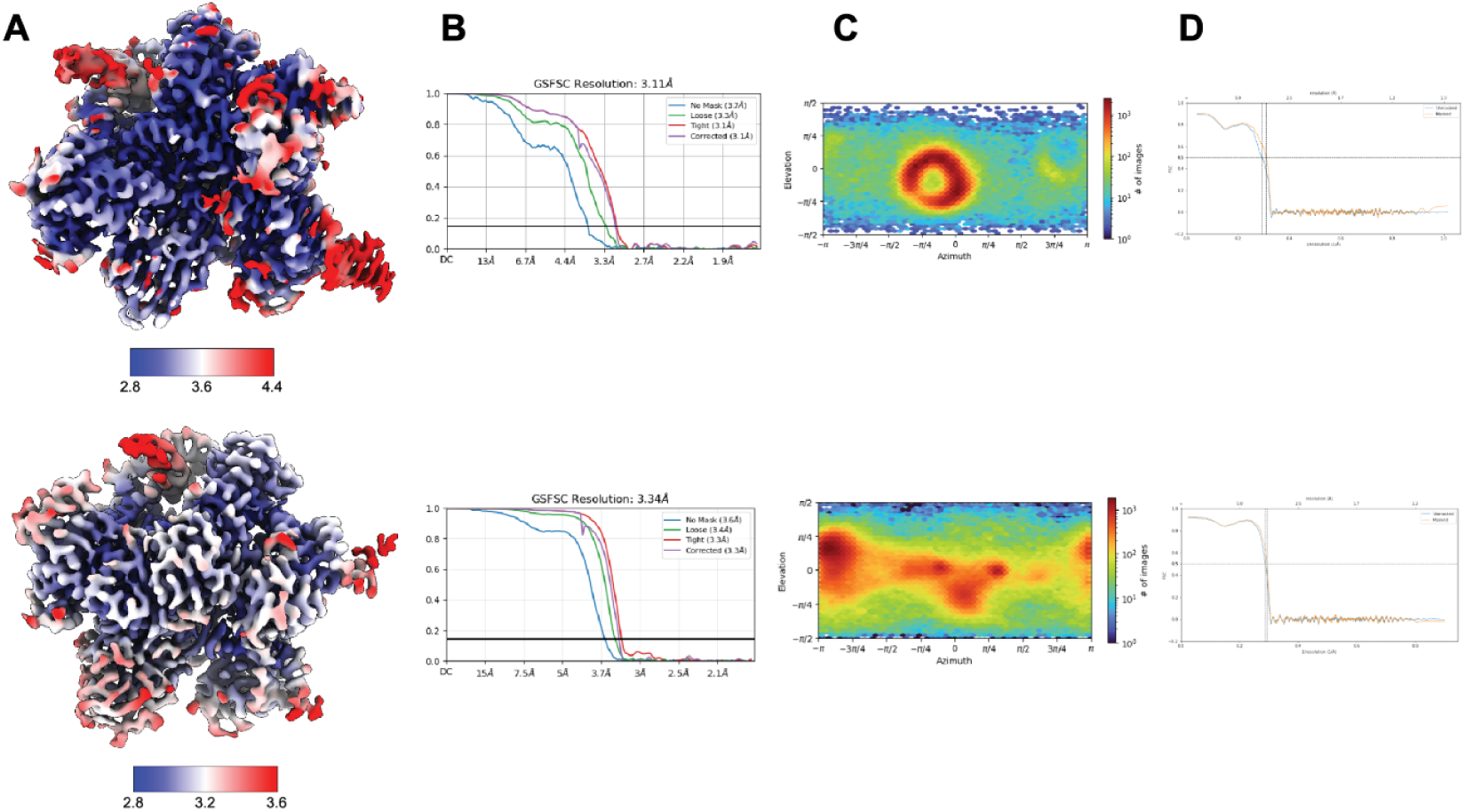
Cas9d cryo-EM data analysis and quality control. **(A)** Sharpened maps colored by local resolution with gold-standard FSC curves using a 0.143 FSC threshold. From top to bottom: 34-29 ternary complex and 34-104 ternary complex. **(B)** All resolutions reported are for sharpened maps at an FSC threshold of 0.143. **(C)** Euler diagrams show the orientation distribution of cryo-EM reconstructions. **(D)** Map to model FSCs are shown for both cryo-EM reconstructions.

**Fig. S8:**
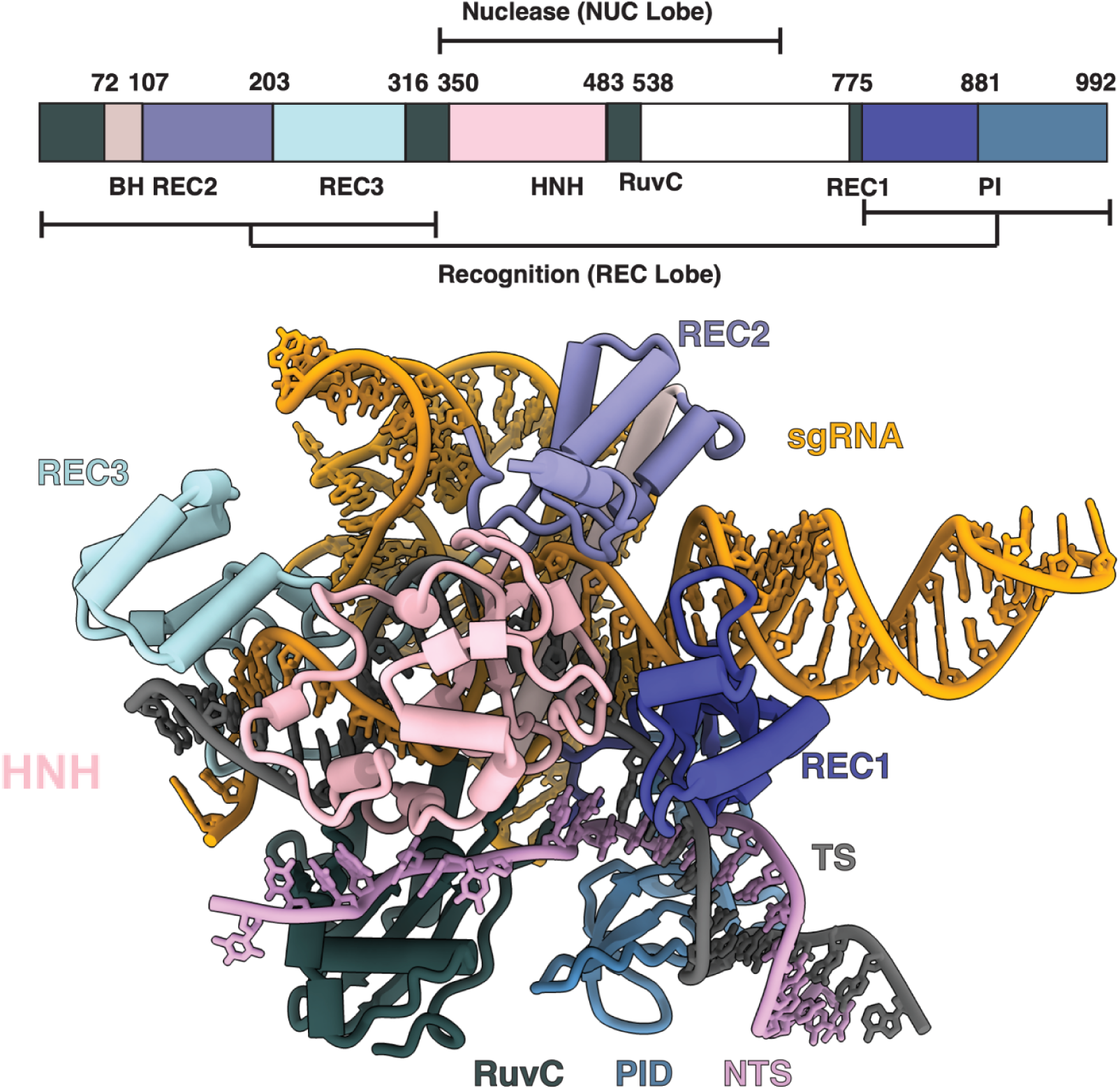
Cryo-EM structure of microABE273. MicroABE273 consists of a monomeric MG68-4 deaminase variant (V83S, D109N, T112R, R154P) inlaid within the RuvC domain (at L516) of MG34-104 (A583R). Top, schematic domain organization of MG34-104. The recognition (REC) lobe comprises REC1 (purple), REC2 (light purple) and REC3 (light blue), while the nuclease lobe contains the HNH (light pink) and RuvC (dark green) domains. The PAM-interacting (PI) domain (royal blue) is located at the C-terminus. Bottom, cryo-EM model of microABE273 bound to sgRNA and target DNA. The sgRNA is shown in orange, the target strand (TS) in grey, and the non-target strand (NTS) in magenta.

## Supplementary Sequences

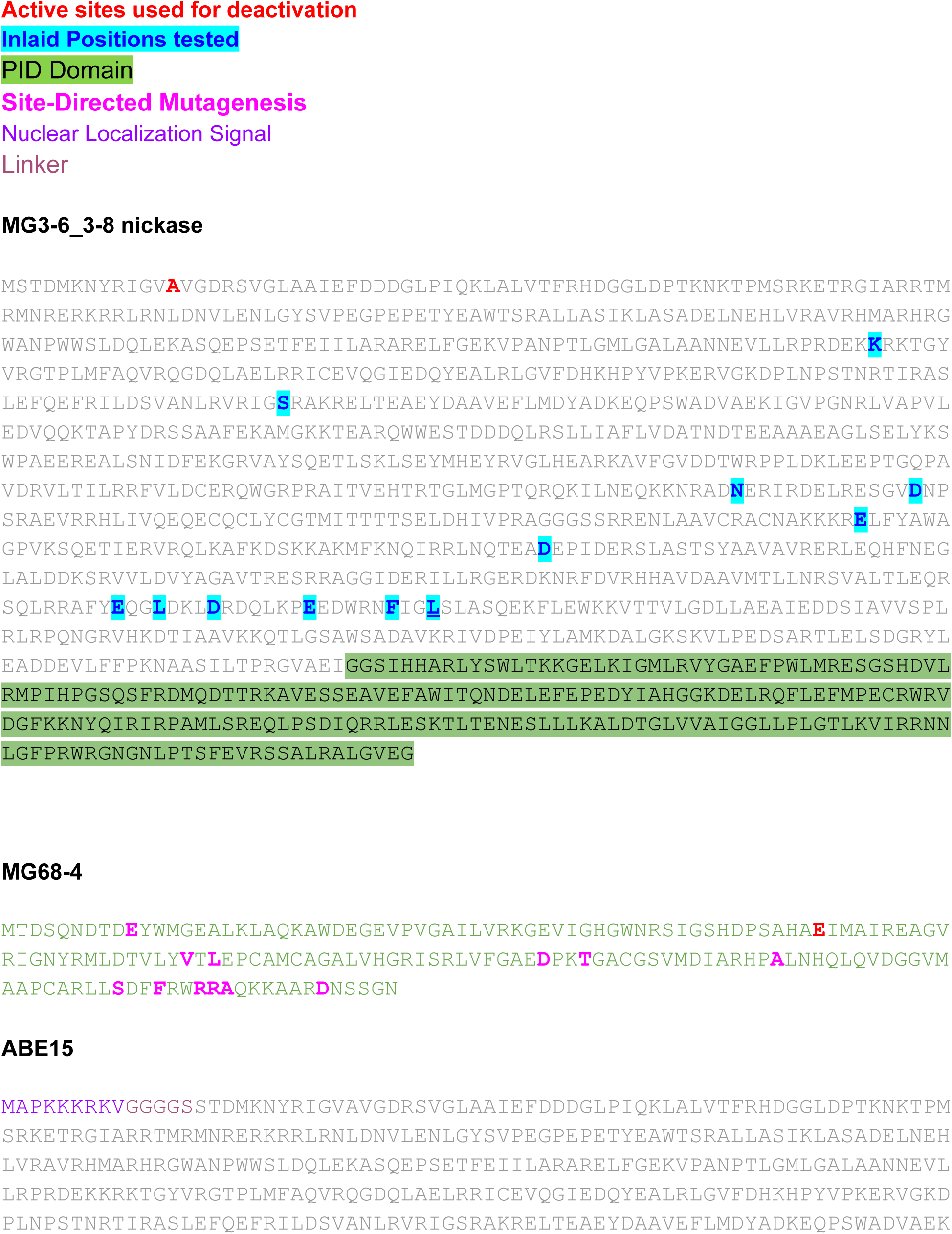
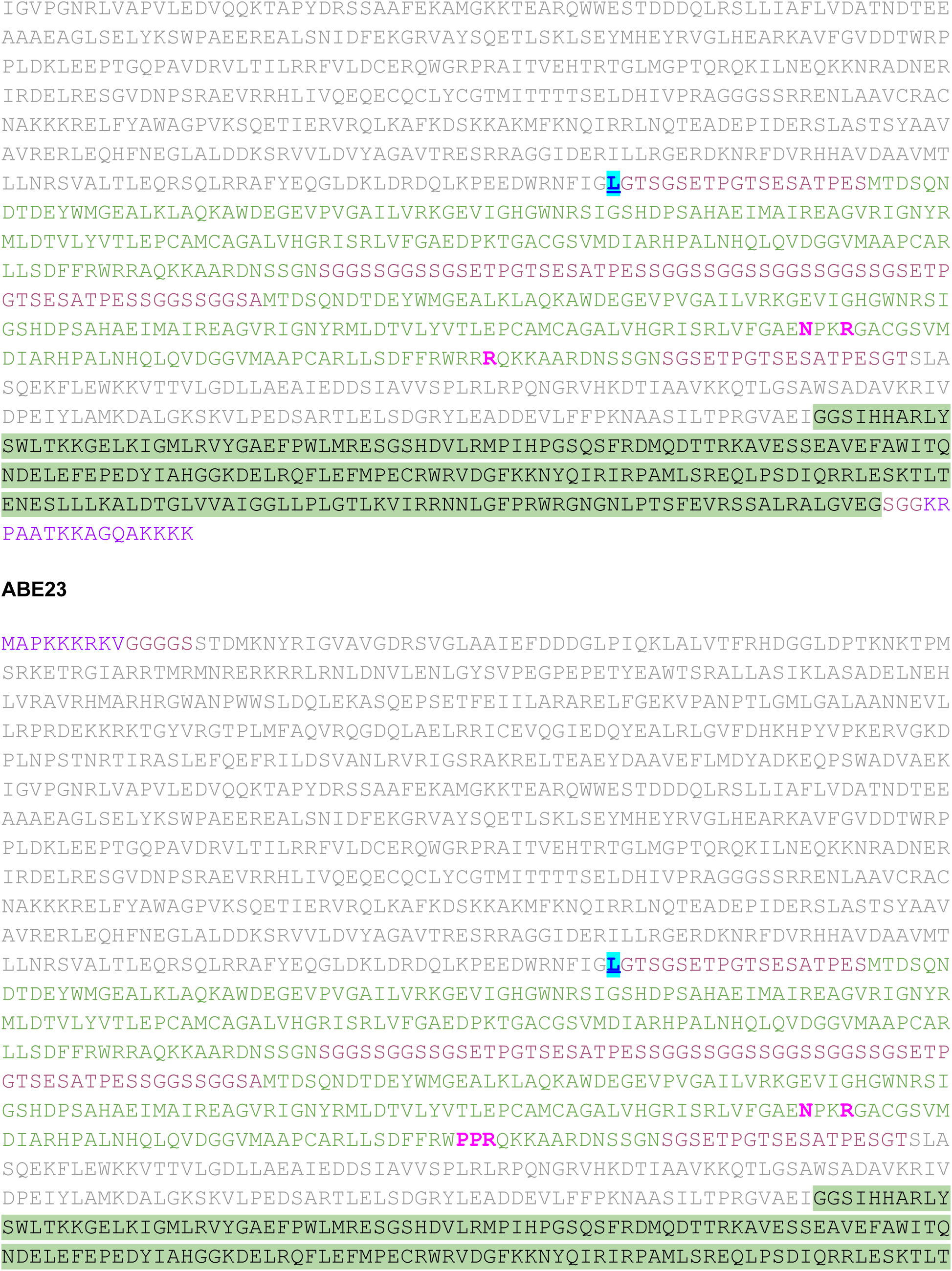
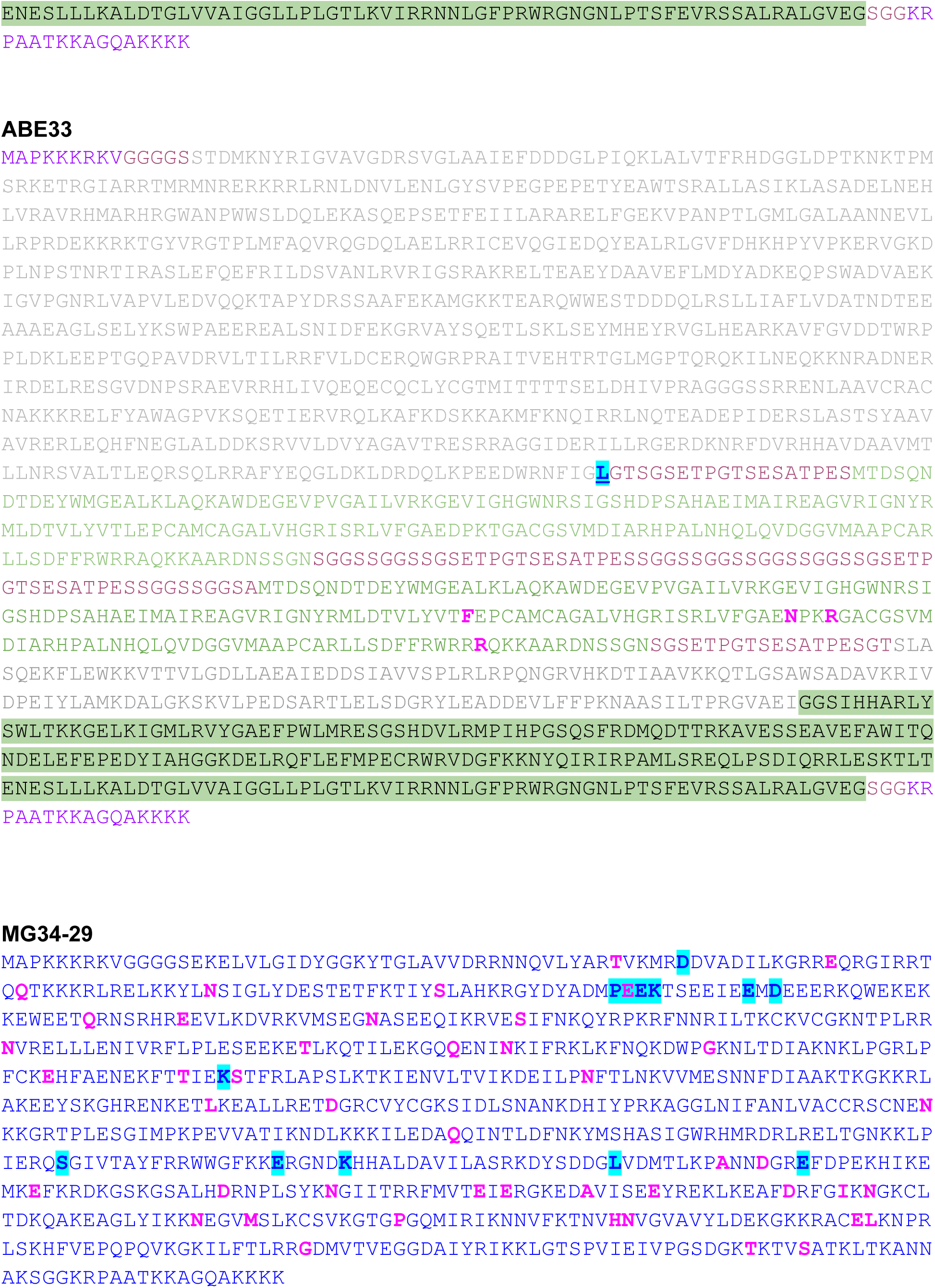
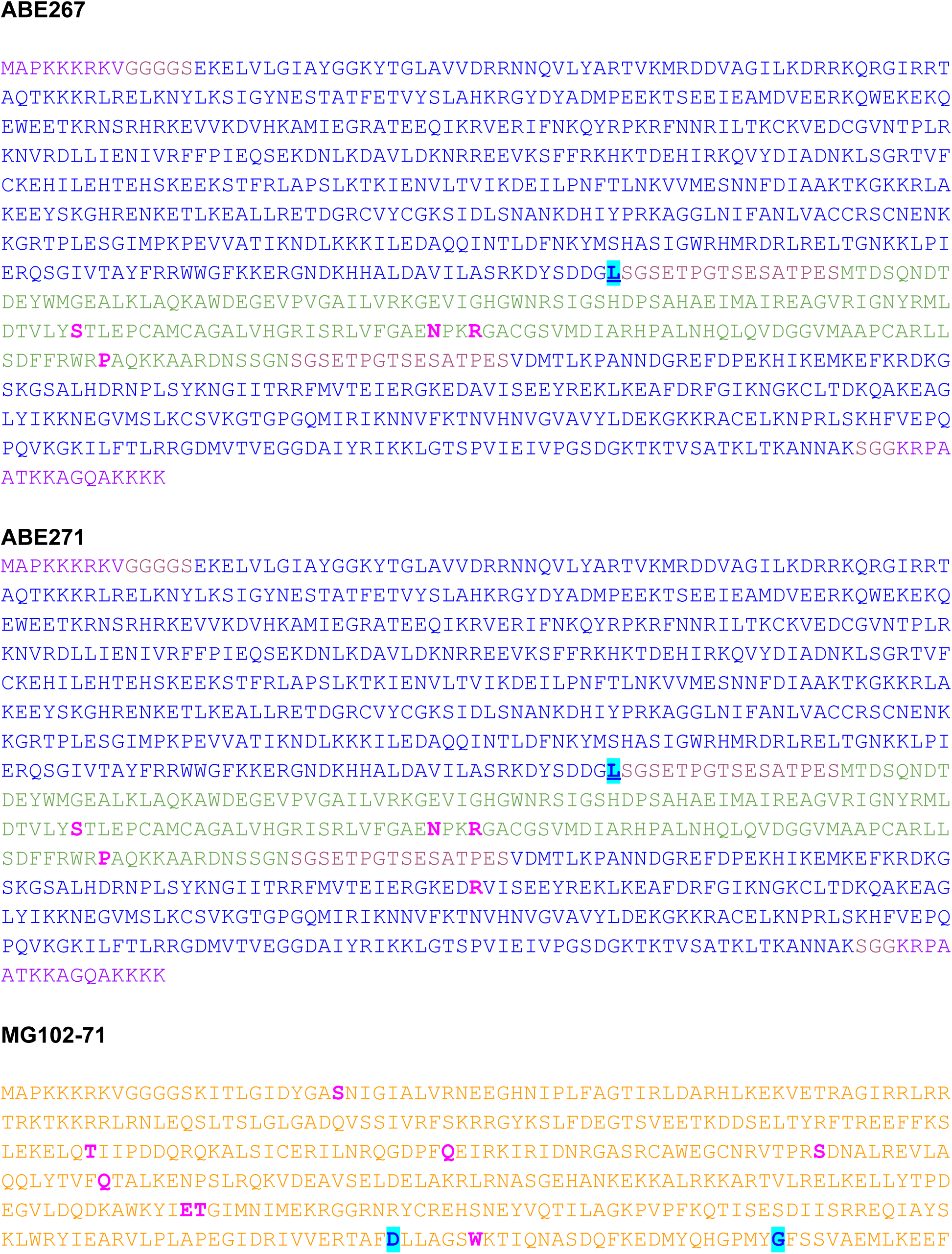
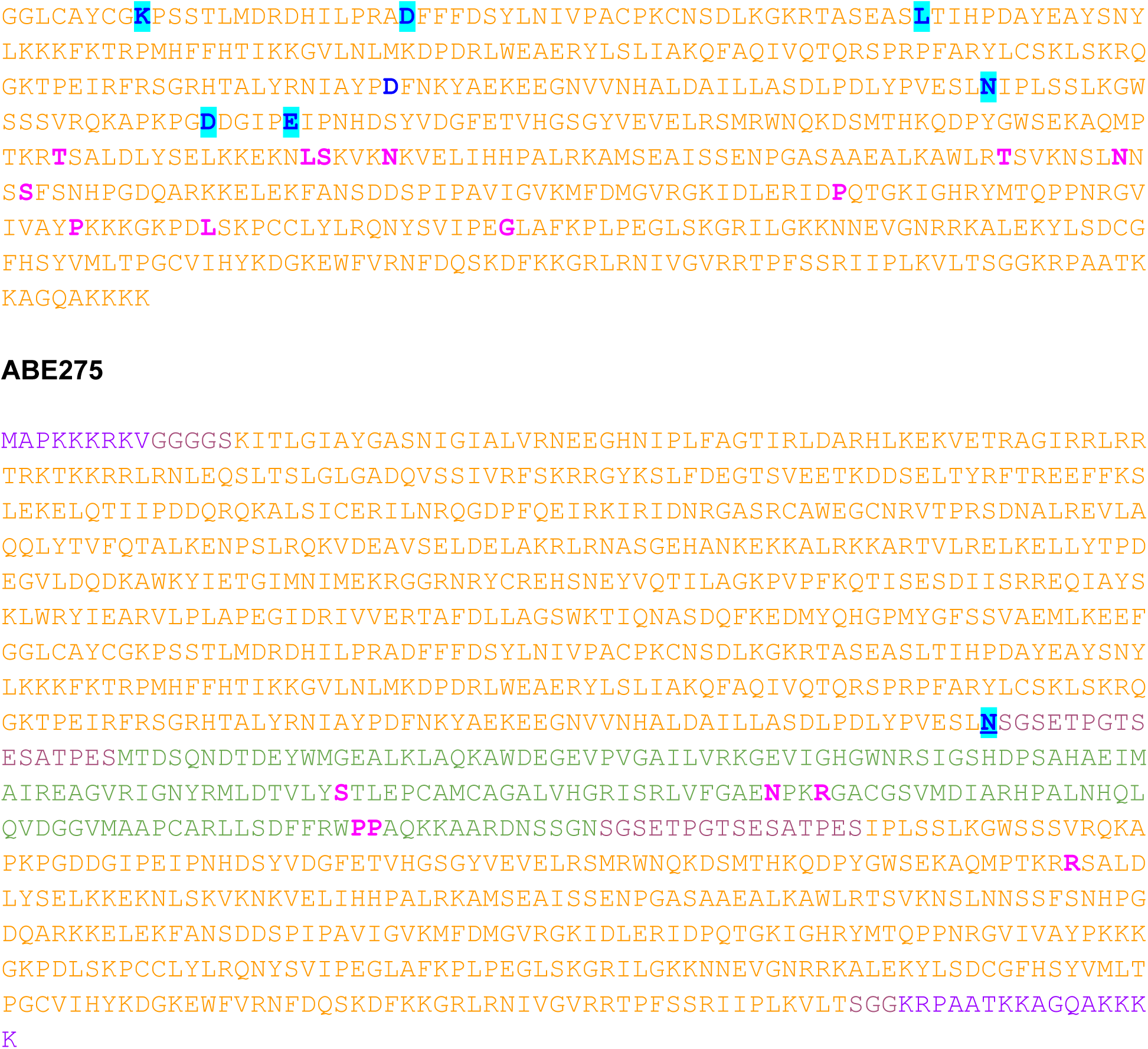

